# DETECTION OF ADAPTIVE SHIFTS ON PHYLOGENIES USING SHIFTED STOCHASTIC PROCESSES ON A TREE

**DOI:** 10.1101/023804

**Authors:** Paul Bastide, Mahendra Mariadassou, Stéphane Robin

**Affiliations:** UMR MIA-Paris, AgroParisTech, INRA, Université Paris-Saclay, 75005, Paris, France; MaIAGE, INRA, Université Paris-Saclay, 78352 Jouy-en-Josas, France

**Keywords:** Random process on tree, Ornstein-Uhlenbeck process, Change-point detection, Adaptive shifts, Phylogeny, Model selection

## Abstract

Comparative and evolutive ecologists are interested in the distribution of quantitative traits among related species. The classical framework for these distributions consists of a random process running along the branches of a phylogenetic tree relating the species. We consider shifts in the process parameters, which reveal fast adaptation to changes of ecological niches. We show that models with shifts are not identifiable in general. Constraining the models to be parsimonious in the number of shifts partially alleviates the problem but several evolutionary scenarios can still provide the same joint distribution for the extant species. We provide a recursive algorithm to enumerate all the equivalent scenarios and to count the effectively different scenarios. We introduce an incomplete-data framework and develop a maximum likelihood estimation procedure based on the EM algorithm. Finally, we propose a model selection procedure, based on the cardinal of effective scenarios, to estimate the number of shifts and prove an oracle inequality.

## 1 Introduction

### 1.1 Motivations: Environmental Shifts

An important goal of comparative and evolutionary biology is to decipher the past evolutionary mechanisms that shaped present day diversity, and more specifically to detect the dramatic changes that occurred in the past (see for instance Losos, 1990; Mahler et al., 2013; Davis et al., 2007; Jaffe et al., 2011). It is well established that related organisms do not evolve independently (Felsenstein, 1985): their shared evolutionary history is well represented by a phylogenetic tree. In order to explain the pattern of traits measured on a set of related species, one needs to take these correlations into account. Indeed, a given species will be more likely to have a similar trait value to her “sister” (a closely related species) than to her “cousin” (a distantly related species), just because of the structure of the tree. On top of that structure, when considering a *functional* trait (i.e. a trait directly linked to the fitness of its bearer), such as shell size for turtles (Jaffe et al., 2011), one needs to take into account the effect of the species environment on its traits. Indeed, a change in the environment for a subset of species, like a move to the Galàpagos Islands for turtles, will affect the observed trait distribution, here with a shift towards giant shell sizes compared to mainland turtles. The observed present-day trait distribution hence contains the footprint of adaptive events, and should allow us to detect unobserved past events, like the migration of one ancestral species to a new environment. Our goal here is to devise a statistical method based on a rigorous maximum likelihood framework to automatically detect the past environmental shifts that shaped the present day trait distribution.

optimum. The value will then be passed down to its descendant, potentially leading to unexpectedly large differences between sub-families of species. The distribution of trait values across extant species hence contains the footprint of adaptive events and should in principle allow us to detect unobserved past events. Our goal here is to devise a statistical framework to automatically detect the past environmental shifts that shaped the present day trait distribution.

### 1.2 Stochastic Process on a tree

We model the evolution of a quantitative adaptive trait using the framework of stochastic processes on a tree. Specifically, given a rooted phylogenetic tree, we assume that the trait evolves according to a given stochastic process on each branch of the tree. At each speciation event, or equivalently node of the tree, one independent copy with the same initial conditions and parameters is created for each daughter species, or outgoing branches.

**Tree Structure**. This model is our null model: it accounts for the tree-induced distribution of trait values in the absence of shifts. Depending on the phenomenon studied, several stochastic processes can be used to capture the dynamic of the trait evolution. In the following, we will use the Brownian Motion (BM) and the Ornstein-Uhlenbeck (OU) processes.

**Figure 1:**
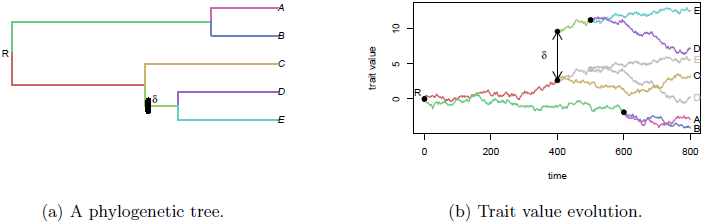
Trait evolution under a Brownian Motion. The ancestral value of the trait is 0, and the observed values (time 800) range from −4 to 11 for extant species. One shift occurs on the parent branch of (D,E), changing the trajectory of their ancestral trait value from the grey one to the colored one. The shift increases the observed dispersion.

**Brownian Motion**. Since the seminal article of Felsenstein (1985), the BM has been used as a neutral model of trait evolution. If (*B_t_*; *t* ≥ 0) is the Brownian motion, a character (*W_t_*;*t* ≥ 0) evolves on a lineage according the the stochastic differential equation: *dW_t_* = *σdB_t_*, *σ*^2^ being a variance parameter. If *μ* is the ancestral value at the root of the tree (*t* = 0), then 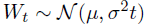. The variance *σ*^2^*t* of the trait is proportional to the time of evolution and the covariance *σ*^2^*t_ij_* between two species *i* and *j* is proportional to their time of shared evolution.

**Ornstein-Uhlenbeck Process.** An unbounded variance is quite unrealistic for adaptive traits (Butler and King, 2004). For that reason, the OU process, that models stabilizing selection around an adaptive optimum (Hansen, 1997) is usually preferred to the BM. It is defined by the stochastic differential equation *dW_t_* = −α(*W_t_* − *β*)*dt* + *σdB_t_*, and has stationary distribution 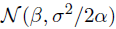. In this equation, *W_t_* is the *secondary optimum* of a species, a trade-off between all selective constraints - e.g. ecological - on the trait and can be approached by the population mean of that species. The term −*α*(*W_t_* − *β*)*dt* of the equation represents the effects of stabilizing selection towards a *primary optimum β*, that depends only on the ecological niche of the species. The selection strength is controlled by the call-back parameter *α*. For interpretation purpose, we will use the *phylogenetic half-life t*_1/2_ = ln(2)/*α*, defined as the time needed for the expected trait value to move half the distance from the ancestral state to the primary optimum (Hansen, 1997). The term *σdB_t_* represents the random effects of uncontrolled factors, ranging from genetic drift to environmental fluctuations. We refer to Hansen (1997); Hansen et al. (2008) for further discussion and deeper biological interpretations of the hypothesis underlying this model of evolution. The aim of our work is to detect environmental shifts.

**Environmental Shifts**. In addition to the previous mechanisms, we assume that abrupt environmental changes affected the ecological niche of some ancestral species. We model these changes as instantaneous shifts in the parameters of the stochastic process. Shifted parameters are inherited along time and thus naturally create clusters of extant species that share the same parameters trajectories. In the BM process, shifts affect the mean value of the trait and are thus instantaneously passed on to the trait itself (see Figure 1) whereas in the OU process, shifts affect the primary optimum *β*. In this case, the trait converges to its new stationary value with an exponential decay of half-life *t_1/2_* inducing a lag that makes recent shifts harder to detect (Hansen and Bartoszek, 2012). In the remainder, we assume that all other parameters (*σ*^2^ for the BM and *σ*^2^, *α* for the OU) are fixed and constant (but see Beaulieu et al., 2012; Rabosky, 2014, for partial relaxations of this hypothesis).

### 1.3 Scope of this article

**State of the Art**. Phylogenetics Comparative Methods (PCM) are an active field that has seen many fruitful developments in the last few years (see Pennell and Harmon, 2013, for an extensive review). Several methods have been specifically developed to study adaptive evolution, starting with the work of Butler and King (2004). Butler and King (2004) only consider shifts in the optimal value *β* whereas Beaulieu et al. (2012) also allow for shifts in the selection strength *α* and the variance *σ*^2^. Both have in common that shift locations are assumed to be known. Several extensions of the model without or with known shifts have also been proposed: Hansen et al. (2008) extended the original work of Hansen (1997) on OU processes to a two-tiered model where *β*(*t*) is itself a stochastic process (either BM or OU). Bartoszek et al. (2012) extended it further to multivariate traits whereas Hansen and Bartoszek (2012) introduced errors in the observations. Expanding upon the BM, (Landis et al., 2013) replaced fixed shifts, known or unknown, by random jump processes using Levy processes. Non-Gaussian models of trait evolution were also recently considered by Hiscott et al. (2015), who adapted Felsenstein’s pruning algorithm for the likelihood computation of these models, using efficient integration techniques. Finally, Ho and Ané (2013) derived consistency results for estimation of the parameters of an OU on a tree and Bartoszek and Sagitov (2015); Sagitov and Bartoszek (2012) computed confidence intervals of the same parameters by assuming an unknown random tree topology and averaging over it.

The first steps toward automatic detection of shifts, which is the problem of interest in this paper, have been done in a Bayesian framework, for both the BM (Eastman et al., 2013) and the OU (Uyeda and Harmon, 2014). Using RJ-MCMC, they provide the user with the posterior distribution of the number and location of shifts on the tree. Convergence is however severely hampered by the size of the search space. The growing use of PCM in fields where large trees are the norm makes maximum likelihood based point estimates of the shift locations more practical. A stepwise selection procedure for the shifts has been proposed in Ingram and Mahler (2013). The procedure adds shifts one at the time and is therefore rather efficient but the selection criterion is heuristic and has no theoretical grounding for that problem, where observations are correlated through the tree structure. These limitations have been pointed out in Ho and Ané (2014b). In this article, the authors describe several identifiability problems that arise when trying to infer the position of the shifts on a tree, and propose a different stepwise algorithm based on a more stringent selection criterion, heuristically inspired by segmentation algorithms.

To rigorously tackle this issue, we introduced a framework where a univariate trait evolves according to an OU process with stationary root state (S) on an Ultrametric tree (U). Furthermore, as the exact position of a shift on a branch is not identifiable for an ultrametric tree, we assume that shifts are concomitant to speciation events and only occur at Nodes (N) of the tree. We refer to this model as OUsun hereafter.

**Our contribution**. In this work, we make several major contributions to the problem at hand. First, we derive a statistical method to find a maximum likelihood estimate of the parameters of the model. When the number of shifts is fixed, we work out an Expectation Maximization (EM) algorithm that takes advantage of the tree structure of the data to efficiently maximize the likelihood. Second, we show that, given the model used and the kind of data available, some evolutionary scenarios remain indistinguishable. Formally, we exhibit some identifiability problems in the location of the shifts, even when their number is fixed, and subsequently give a precise characterization of the space of models that can be inferred from the data on extant species. Third, we provide a rigorous model selection criterion to choose the number of shifts needed to best explain the data. Thanks to our knowledge of the structure of the spaces of models, acquired through our identifiability study, we are able to mathematically derive a penalization term, together with an oracle inequality on the estimator found. Fourth and finally, we implement the method on the statistical software **R** (R Core Team, 2014), and show that it correctly recovers the structure of the model on simulated datasets. When applied to a biological example, it gives results that are easily interpretable, and coherent with previously developed methods. All the code used in this article is publicly available on GitHub (https://github.com/pbastide/PhylogeneticEM).

**Outline**. In Section 2, we present the model, using two different mathematical point of views, that are both useful in different aspects of the inference. In Section 3, we tackle the identifiability problems associated with this model, and describe efficient algorithms to enumerate, first, all equivalent models within a class, and, second, the number of truly different models for a given number of shifts. These two sections form the foundation of Section 4, in which we describe our fully integrated maximum likelihood inference procedure. Finally, in Sections 5 and 6, we conduct some numerical experiments on simulated and biological datasets.

## 2 Statistical Modeling

### 2.1 Probabilistic Model

**Figure 2:.**
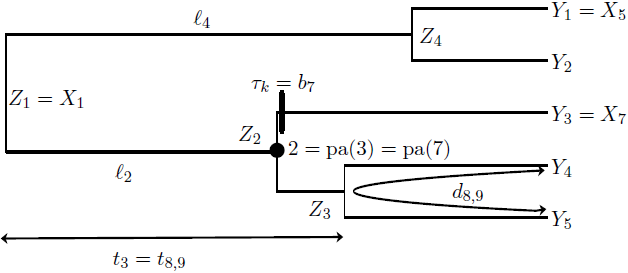
A rooted and time calibrated phylogenetic tree with the notations used to parametrize the tree (*l*,*t*,*d*,*b*) and the observed (*Y*) and non-observed (*Z*) variables.

**Tree Parametrization.** As shown in Figure 2, we consider a rooted tree 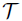 with *n* tips and *m* internal nodes (*m* = *n* − 1 for binary trees). The internal nodes are numbered from 1 (the root) to *m*, and the tips from *m* + 1 to *m* + *n*. Let *i* be an integer, 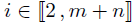. Then pa(*i*) is the unique parent of node *i*. The branch leading to *i* from pa(*i*) is noted *b_i_* and has length *ℓ*_*i*_ = *t_i_* − *t*_pa_(_*i*_) where *t_i_* is the time elapsed between the root and node *i*. By convention, we set *t*_1_ = 0 and *t*_pa(1)_ = − ∞ for the root. The last convention ensures that the trait follows the stationary distribution (if any) of the process at the root. We denote Anc(*i*) = {pa^r^(*i*): *r* ≥ 0} the set composed of node *i* and of all its ancestors up to the root. For a couple of integers (*i*,*j*), 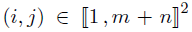, nodes *i* and *j* are at phylogenetic distance *d_ij_* and the time of their most recent common ancestor (mrca) is *t_ij_*. We consider ultrametric trees, for which *t_m+1_* = …= *t_*m*+*n*_* =: *h* and note *h* the tree height. In the following, the tree is fixed and assumed to be known.

**Trait Values**. We denote by **X** the vector of size *m* + *n* of the trait values at the nodes of the tree. We split this vector between non-observed values **Z** (size *m*) at the internal nodes, and observed values **Y** (size *n*) at the tips, so that **X**^*T*^ = (**Z**^*T*^, **Y**^*T*^). According to our model of trait evolution, the random variable 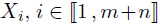, is the result of a stochastic process stopped at time *t_i_*. In the following, we assume that the inference in the BM case is done conditionally to a fixed root value *X*_1_ = *μ*. In the OUsun case, we assume that the root trait value is randomly drawn from the stationary distribution: 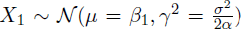, where *β*_1_ is the ancestral optimal value.

**Shifts.** We assume that *K* shifts occur on the tree, 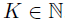. The *k*^th^ shift, 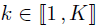, occurs at the beginning of branch τ_k_, 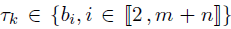, and has intensity *δ_k_*, *δ_k_* ∈ ℝ. The interpretation of this intensity depends on the process. In the following, we use the vector Δ of shifts on the branches, of size *m* + *n*, with *K* + 1 non-zero entries, and defined as follows (see example 2.1):

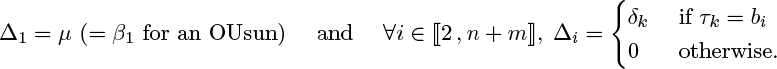

Note that no proper shift occurs on the root branch, but that the root trait value or mean, *μ*, is formalized as an initial fictive shift on this fictive branch.

**Parameters.** The parameters needed to describe an OUsun (respectively, a BM) are *θ* = (γ, *α*,**Δ**) (resp. *θ* = (*σ*, **Δ**)). Note that, as *σ*^2^ = 2*αγ*^2^, only the two parameters *α* and γ are needed to describe the OUsun. We denote by OUsun(***θ***) (resp. BM(6)) the OUsun (resp. BM) process running on the tree with parameters ***θ***.

### 2.2 Incomplete Data Model Point of View

If the trait values were observed at all nodes of the tree, including ancestral ones, shifts would be characterized by unexpectedly large differences between a node and its parent. A way to mimic this favorable case is to use an incomplete data model, as described below. This representation of the model will be useful for the parametric inference using an EM algorithm (Section 4.1).

**Brownian Motion**. As the shifts occur directly in the mean of the process, we get: 
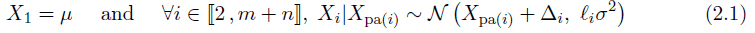

The trait value at node *i*, 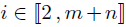, is centered on the value of its parent node *X*_pa_(_*i*_), with a variance proportional to the evolution time *ℓ_i_* between *i* and pa(*i*). The effect of a non-zero shift Δ_*i*_ on branch *b_i_* is simply to translate the trait value by Δ_*i*_.

**Ornstein-Uhlenbeck**. The shifts occur on the primary optimum *β*, which is piecewise constant. As the shifts are assumed to occur at nodes, the primary optimum is entirely defined by its initial value *β_1_* and its values *β*_2_,…, *β*_*n*+*m*_ on branches of the tree, where *β_i_* is the value on branch *b_i_* leading to node *i*.

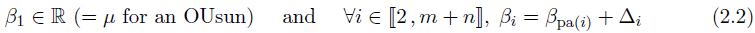

Assuming that the root node is in the stationary state, we get:

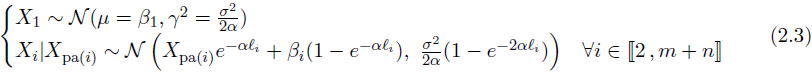

The trait value at node *i* depends on both the trait value at the father node *X*_pa(*i*)_ and the value *β_i_* of the primary optimum on branch *b_i_*. Contrary to the BM case, the shifts only appear indirectly in the distributions of *X_i_*s, through the values of *β*, and with a shrinkage of 1 − *e*^−*αd*^ for shifts of age *d*, which makes recent shifts (*d* small compared to 1/*α*) harder to detect.

### 2.3 Linear Regression Model Point of View

A more compact and direct representation of the model is to use the tree incidence matrix to link linearly the observed values (at the tips) with the shift values, as explained below. We will use this linear regression framework for the Lasso (Tibshirani, 1996) initialization of the EM (Section 4.1) and the model selection procedure (Section 4.2). It will also help us to explore identifiability issues raised in the next section.

**Matrix of a tree.** It follows from the recursive definition of **X** that it is a Gaussian vector. In order to express its mean vector given the shifts, we introduce the tree squared matrix **U**, of size (*m* + *n*), defined by its general term: 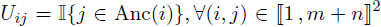. In other words, the *j*^th^ column of this matrix, 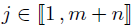, is the indicator vector of the descendants of node *j*. To express the mean vector of the observed values **Y**, we also need the sub-matrix **T**, of size *n* × (*m* + *n*), composed of the bottom n rows of matrix **U**, corresponding to the tips (see example 2.1 below). Likewise, the *i*^th^ row of **T**, 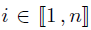, is the indicator vector of the ancestors of leaf *m* + *i*.

**Brownian Motion**. From the tree structure, we get:

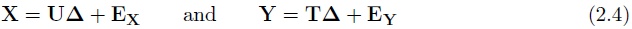

Here, 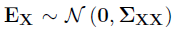 is a Gaussian error vector with co-variances [∑_*XX*_]_*ij*_ = *σ*^2^*t_ij_* for any 1 ≤ *i*,*j* ≤ *m* + *n*, and **E**_**Y**_ is the vector made of the last *n* coordinates of **E_x_**.

**Ornstein-Uhlenbeck**. For the OUsun, shifts occur on the primary optimum, and there is a lag term, so that: 
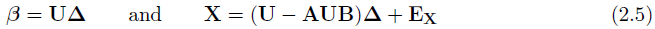
where 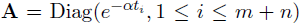 and 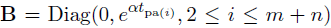 are diagonal matrices of size *m* + *n*. As previously, 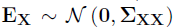, but 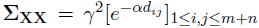. As the tree is ultrametric, this expression simplifies to the following one when considering only observed values: 
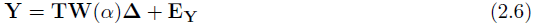
where **E**_**y**_ is the Gaussian vector made of the last *n* coordinates of **E**_**X**_, and 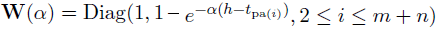 is a diagonal matrix of size *m* + *n*. Note that if *α* is positive, then *α*(*h* − *t*_pa(*i*)_ > 0 for any 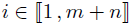, and **W**(*α*) is invertible.

#### *Example* 2.1.

The tree presented in Figure 2 has five tips and one shift on branch 4 + 3 = 7, so:

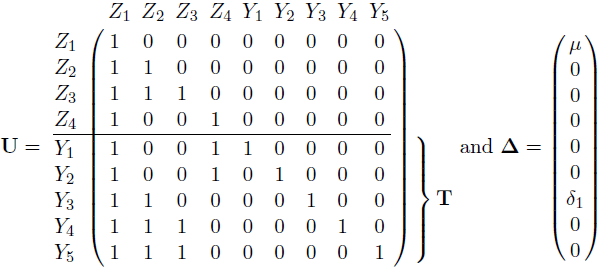

And, respectively, for a BM or an OUsun (*μ* =*β*_1_): 
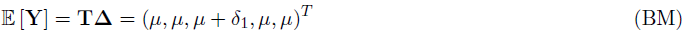

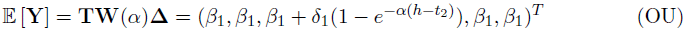

**Space of Expectations**. Expressions 2.4 and 2.6 allow us to link the parameter ***θ*** to the probability distribution of observations ***Y*** and to explore identifiability issues. In this linear formulation, detecting shifts boils down to identifying the non-zero components of **Δ**. The following lemma highlights the parallels between solutions of the BM and OUsun processes:

#### Lemma 2.1

(Similar Solutions). *Let* **m_Y_** ∈ ℝ^*n*^ *be a vector*, 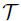 *an ultrametric tree*, *α a positive real number*, and *σ*, *γ non-negative real numbers. Then there exists at least one vector* **Δ**^BM^, **Δ**^*BM*^ ∈ ℝ^*m*+*n*^ *(respectively, **Δ**^*OU*^ ∈ ℝ^*m*+*n*^)*, *such that the vector of expectations at the tips of a BM*(*σ*, **Δ**^*BM*^) *(respectively, an OUsun*(γ,α, **Δ**^*OU*^)) *running on the tree* 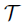 is exactly **m**_**Y**_.

*Furthermore*, **Δ**^*BM*^ *is a solution to this problem for the *BM* if and only if* **Δ**^*OU*^ = **W**(*α*)^−1^ **Δ**^BM^ *is a solution for the OUsun*, *and* **Δ**^*BM*^ *and* **W**(*α*)^−1^**Δ**^*BM*^ *have the same support*. *These two vectors are said to be similar*.

#### *Proof*.

The first part of this lemma follows directly from formulas 2.4 (BM) and 2.6 (OU). Indeed, the maps **Δ** ↦ **TΔ** and **Δ** ↦ **TW**(*α*)**Δ** both span ℝ^*n*^. The second part of the lemma is a consequence of **W**(*α*) being diagonal and invertible (for *α* > 0).

#### *Remark* 2.1.

Lemma 2.1 shows that the OUsun and BM processes that induce a given **m**_**Y**_ use shifts located on the same branches, although they may differ on other parameters.

## 3 Identifiability and Complexity of a Model

### 3.1 Identifiability Issues

As we only have access to **Y**, and not **X**, we only have partial information about the shifts occurrence on the tree. In fact, several different allocations of the shifts can produce the same trait distribution at the tips, and hence are not identifiable. In other words, there exists parameters *θ* ≠ *θ′* with the same likelihood function: *p_θ_*(·) = *p_θ′_*(·). Note that the notion of identifiability is intrinsic to the model and affects all estimation methods. Restricting ourselves to the parsimonious allocations of shifts only partially alleviates this issue, and, using a “random cluster model” representation of the problem, we are able to enumerate, first, all the equivalent solutions to a given problem, and, second, all the equivalence classes for a given number of shifts.

**Figure 3:**
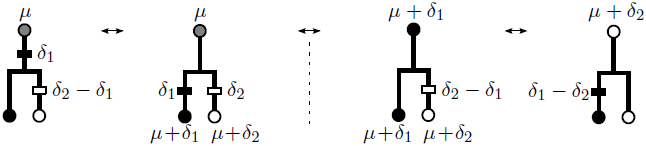
Equivalent allocations in the BM case. Mean tip values are represented by colors and equal for all allocations. The two allocations on the right are parsimonious.

**No Homoplasy Assumption**. We assume in the following that there is no convergent evolution. This means that each shift creates a new (and unique) mean trait value for extant species that are below it. This assumption is reasonable considering that shifts are real valued and makes the model similar to “infinite alleles” models in population genetics. This assumption confines but does not eliminate the identifiability issue, as seen in Figure 3.

#### 3.1.1 Definition of the problem

Figure 3 shows a simple example where the model is not identifiable in the BM case. Here, four distinct allocations give the same mean values (*μ* + *δ_1_*,*μ* + *δ*_2_) at the tips. The lack of identifiability is due to the non-invertibility of the tree matrix **T**.

##### Proposition 3.1

(Kernel of the Tree Matrix **T**). *Let i be an internal node, 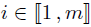, *with L_i_ children nodes* 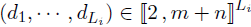. Then the vector* **K**^*i*^ *defined as follow*: 
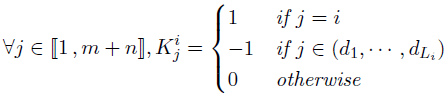
*is in the kernel of* **T**. *In addition, the m vectors constructed this way form a basis of the kernel space of* **T**.

These kernel vectors effectively “cancel out” a shift on a branch by balancing it with the opposite shift on all immediate child branches. Note that the root mean value is treated as a shift. The following lemma describes the relationships that exist between these kernel vectors and the tree matrix U defined in Section 2.3. The technical proofs of Proposition 3.1 and Lemma 3.1 are postponed to Appendix A.

##### Lemma 3.1.

*Let b be the canonical basis of* ℝ^*m*+*n*^, *and S a supplementary space of* ker(**T**). *Then b*′ = (**K**^1^, …, **K**^*m*^, **b**_*m*+1_, …, **b**_*m*+*n*_) *is a basis adapted to the decomposition* ker(**T**) ⊕ *S*, *and the matrix* **U** *(as defined in Section 2.3) is the change of basis matrix between b and b′*. *As a consequence*, **U** *is invertible*.

**“Random Cluster Model” Representation**. When inferring the shifts, we have to keep in mind this problem of non-identifiability, and be able to choose, if necessary, one or several possible allocations among all the equivalent ones. In order to study the properties of the allocations, we use a *random cluster model*, as defined in Mossel and Steel (2004). The following definition states the problem as a node coloring problem.

##### Definition 3.1

(Node Coloring). Let 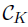 be a set of *K* arbitrary “colors”, *K* ∈ ℕ*. For a given shift allocation, the color of each node is given by the application 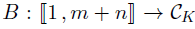 recursively defined in the following way:

- Choose a color 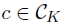 for the root: *B*(1) = *c*.
- For a node *i*, 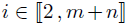, set *B*(*i*) to *B*(pa(*i*)) if there is no shift on branch *i*, otherwise choose another color *c*, 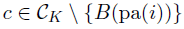, and set *B*(*i*) to *c*.

Hereafter, we identify 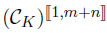 with 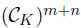 and refer to a node coloring indifferently as an application or a vector.

As the shifts only affect 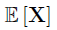 and we only have access to 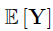, we identify colors with the distinct values of 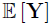:

##### Definition 3.2

(Adapted Node Coloring). A node coloring is said to be *adapted* to a shifted random process on a tree if two *tips* have the same color if and only if they have the same mean value under that process.

##### Proposition 3.2

(Adapted Coloring for BM and OUsun). *Let σ and* γ *be two non-negative real numbers, and α a positive real number. Then*:

i. *In the BM case, if 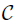 is the set of possible mean values taken by the nodes of the tree, then the knowledge of the node colors is equivalent to the knowledge of* **Δ**. *Furthermore, the associated node coloring is adapted to the original BM*.
ii. *In the OUsun case, from lemma 2.1, we can find a similar BM process, i.e. with shifts on the same branches. Then the knowledge of the node coloring associated to this similar BM process is equivalent to the knowledge of the vector of shifts of the OUsun, and the node coloring obtained is adapted to the original OUsun*.

##### *Proof of Proposition 3.2*.

The proof of (*i*) relies on expression 2.4, that states that 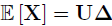. Defining 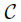 as the set off all distinct values of 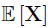, we can identify 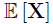 with the node coloring application that maps any node *i* with 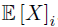. Since **U** is invertible (see lemma 3.1 above), we can go from one formalism to the other.

For (*ii*), we use lemma 2.1 to find a similar BM, and then use (i).

From now on, we will study the problem of shifts allocation as a discrete-state coloring problem.

#### 3.1.2 Parsimony

As we saw on Figure 3 there are multiple colorings of the internal nodes that lead to a given tips coloring. Among all these solutions, we choose to study only the *parsimonious* ones. This property can be seen as an optimality condition, as defined below:

##### Definition 3.3

(Parsimonious Allocation). Given a vector of mean values at the tips produced by a given shifted stochastic process running on the tree, an adapted node coloring is said to be *parsimonious* if it has a minimum number of color changes. We denote by 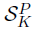 the set of parsimonious allocations of *K* shifts on the (*m* + *n* − 1) branches of the tree (not counting the root branch).

As *K* shifts cannot produce more than *K* + 1 colors, we can define an application 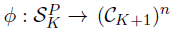 that maps a parsimonious allocation of shifts to its associated tip partition.

##### Definition 3.4

(Equivalence). Two allocations are said to be *equivalent* (noted ~) if they produce the same partition of the tips and are both parsimonious. Mathematically: 
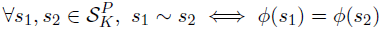

In other words, two allocations are equivalent if they produce the same tip *coloring* up to a permutation of the colors. Given 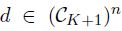 a coloring of the tips of 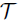 with *K* + 1 colors, *ϕ*^−1^(*d*) is the set of equivalent parsimonious node coloring that coincide with *d* (up to a permutation of the colors) on the tree leaves.

Several dynamic programming algorithms already exist to compute the minimal number of shifts required to produce a given tips coloring, and to find one associated parsimonious solution (see Fitch, 1971; Sankoff, 1975; Felsenstein, 2004). Here, we need to be a little more precise, as we want to both count and enumerate all possible equivalent node colorings associated with a tip coloring. For the sake of brevity, we only present the algorithm that counts |*ϕ*^−1^(*d*)|, for 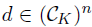. This algorithm can be seen as a corollary of the enumeration algorithm (presented and proved in Appendix B) and an extension of Fitch algorithm where we keep track of both the cost of an optimal coloring and the number of such colorings. It has *O*(*K*^2^*Ln*) time complexity where *L* is the maximal number of children of the nodes of the tree.

##### Proposition 3.3

(Size of an equivalence class). *Let d be a coloring of the tips*,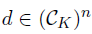, *and let i be a node of tree* 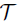 with *L_*i*_ daughter nodes* (*i*_1_, …, *i*_*L_i_*_), *L*_*i*_ ≥ 2. *Denote by* 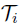 *the sub-tree rooted at node i*.

*For* 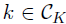, *S*_*i*_(*k*) *is the cost of starting from node i with color k*, *i.e. the minimal number of shifts needed to get the coloring of the tips of* 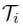 *defined by d*, *when starting with node i in color k*. *Denote by T_i_*(*k*) *the number of allocations on 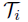 that achieve cost S_i_*(*k*).

*If i is a tip* (*m* + 1 ≤ *i* ≤ *m* + *n*), *then*, 
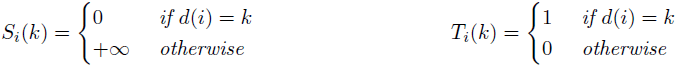

*Otherwise*, *if i is a node*, *for* 1 ≤ *l* ≤ *L_i_*, *define the set of admissible colors for daughter i_l_*: 
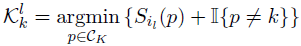

*As these sets are not empty*, *let* 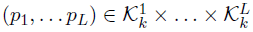. *Then*: 
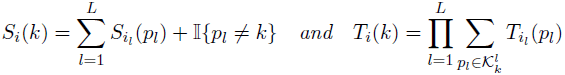

*At the root*, *if* 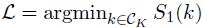, *then* 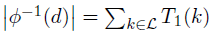.

**OU Practical Case**. We can illustrate this notion on a simple example. We consider an OUsun on a random tree of unit height (total height *h* = 1). We put three shifts on the tree, producing a given trait distribution. Then, using proposition 3.2 and our enumeration algorithm, we can reconstruct the 5 possible allocations of shifts that produce the exact same distribution at the tips. These solutions are shown in Figure 4. Note that the colors are not defined by the values of the optimal regime *β*, but by the mean values 𝔼 [**Y**] of the process at the tips. As a result, the groups shown in blue and red in the first solution have the same optimal value in this configuration, but not in any other. The second solution shown illustrates the fact that all the shifts values are inter-dependent, as changing the position of only one of them can have repercussions on all the others. Finally, the third solution shows that the timing of shifts matters: to have the same impact as an old shift, a recent one must have a much higher intensity (under constant selection strength such as in the OUsun). preferring cases with lower shifts values. According to this rule, solutions 2 or 4 would be preferred.

**Figure 4:**
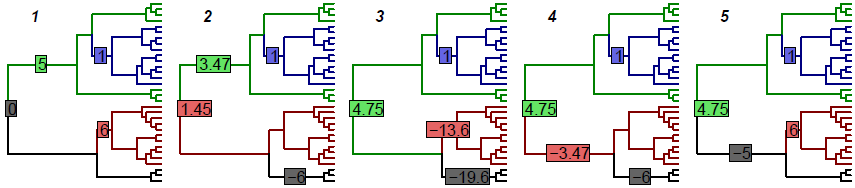
Five equivalent shift allocations that produce colorings that are adapted to an OUsun, with *α* = 3 and *γ*^2^ = 0.1. The box at the root represents the ancestral optimum *β*_1_, and the boxes on the branches represent the positions and values of the shifts on the optimal value. While accounting for very different evolutionary scenarios, all allocations produce the same trait distribution at the tips.

**Possible relaxation of the No-homoplasy assumption**. Note that the algorithms used for counting and enumerating the configurations of an equivalence class are valid even without the no-homoplasy hypothesis. The no-homoplasy hypothesis is however crucial in the next Section to establish a link between the number of shifts and the number of distinct tips colors.

#### 3.2 Complexity of a Collection of Models

**Number of different tips colors**. As we make the inference on the parameters with a fixed number of shifts *K* (see Section 4.1), we need a model selection procedure to choose *K*. This procedure depends on the *complexity* of the collection of models that use *K* shifts, defined as the number of *distinct* models. To do that, we count the number of *tree-compatible* partitions of the tips into *K* + 1 groups, as defined in the next proposition:

##### Proposition 3.4.

*Under the no homoplasy assumption, an allocation of *K* shifts on a tree is parsimonious if and only if it creates exactly K* + 1 *tip colors*. *The tip partition into K* + 1 *groups associated with this coloring is said to be* tree-compatible. *The set* 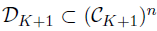 *of such partitions is the image of 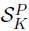 by the map *ϕ* defined in the previous section*.

##### *Proof of Proposition 3.4*.

First, note that *K* shifts create at most *K* + 1 colors. If each shift produces a new tip mean value (no homoplasy), the only way to create *K* or less colors is to “forget” one of the shifts, i.e. to put shifts on every descendant of the branch where it happens. Such an allocation is not parsimonious, as we could just add the value of the forgotten shift to all its descendant to get the same coloring of the tips with one less shift. So a parsimonious allocation cannot create less than *K* + 1 colors, and hence creates exactly *K* + 1 colors.

Reciprocally, if an allocation with *K* shifts that produces *p* groups is not parsimonious, then we can find another parsimonious one that produces the same *p* groups with *p* − 1 shifts, with *p* − 1 < *K*, i.e. *p* < *K* + 1. So, by contraposition, if the allocation produces *K* + 1 groups, then it is parsimonious.

Using the equivalence relation defined in Definition 3.4, we can formally take the quotient set of 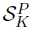 by the relation ~ to get the set of parsimonious allocations of *K* shifts on the *m* + *n* − 1 branches of the tree that are identifiable: 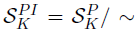. In other words, the set 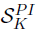 is constituted of one representative of each equivalence class. Under the no homoplasy assumption, there is thus a bijection between identifiable parsimonious allocations of *K* shifts and tree-compatible partitions of the tips in *K* + 1 groups: 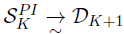.

The number 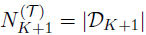 is the complexity of the class of models with *K* shifts defined as the number of distinct identifiable parsimonious possible configurations one can get with *K* shifts on the tree. To compute 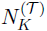, we will need 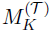 the number of *marked* tree-compatible partitions in *K* groups. These are composed of all the tree-compatible partitions where one group, among those that could be in the same state as the root, is distinguished with a mark (see example 3.1 below).

##### *Example* 3.1

(Difference between 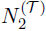 and 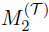.

- If we consider only unmarked partitions, then colorings 1, 2 and 3 induce the same partitions as, respectively, colorings 4, 5 and 6, and 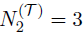.
- For marked partitions, fix the root state to an arbitrary color, for instance white, and consider the white group as marked. Then colorings 5 and 6 are not tree-compatible (they require two shifts). And although they induce the same partition, colorings 1 and 4 correspond to different marked partitions: each marks a different group of leaves. Therefore 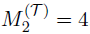.

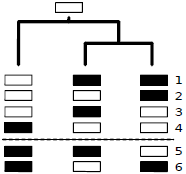
Partitions in 2 groups.

##### Proposition 3.5

(Computation of the Number of Equivalent Classes). *Let i be a node of tree* 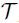, and *K* ∈ ℕ*.

*If i is a tip*, *then* 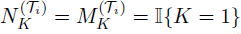.

*Else*, *if i is a node with *L*_i_ daughter nodes* (*i*_1_, …, *i*_*L_i_*_), *L_i_* ≥ 2, *then*: 
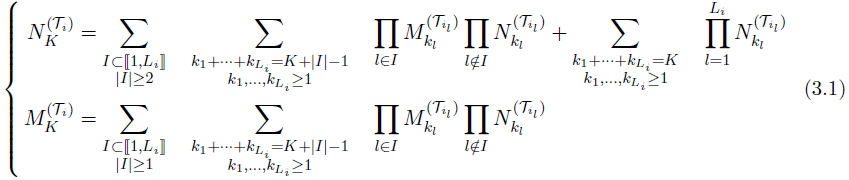

*In the binary case*, *this relation becomes*, *if i has two daughters i*_ℓ_ *and i*_*r*_: 
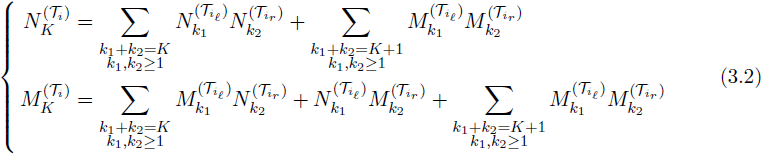

##### *Proof*.

We will prove this proposition in the binary case, the general case being a natural extension of it. If 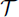 is a binary tree with 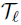 and 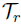 as left and right sub-trees, one faces two situations when partitioning the tips in *K* groups:

- The left and right sub-trees have no group in common. Then, the number of groups in 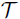 is equal to the number of groups in its two sub-trees, and there are 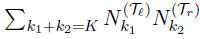 such partitions. This is the first term of the equation on 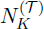 in 3.2.
- The left and right sub-trees have at least one group in common. Then, from the no homoplasy assumption, they have exactly one group in common: the ancestral state of the root. Suppose that this ancestral state is marked. Then it must be present in the two sub-trees, and there are 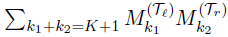 such partitions. This ends the proof of the formula on 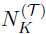.

To get the formula on 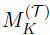, we use the same kind of arguments. The second part of the formula is the same as the one for 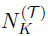, and the first part corresponds to trees for which the marked partition is present in only one of the two sub-trees.

The complexity of the algorithm described above is *O*(2^*L*^(*K* + *L*)^*L*^*Ln*). Note that 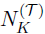 depends on the topology of the tree 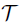 in general. However, if the tree is binary, a closed form solution of the recurrence relation 3.1, which does not depend on the topology, exists.

##### Corollary 3.1

(Closed Formula Binary Trees). *For a rooted binary tree with n tips*, *we have:* 
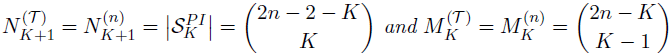

The demonstration of this formula is not straightforward, and is based on a Vandermonde-like equality, detailed in Appendix C. The formula is then obtained using a strong induction on the number of tips of the tree.

##### *Remark* 3.1.

Note that, when *K* is large compared to 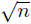, the average number of configurations per equivalence class goes to infinity. This can be checked by comparing the total number of configurations 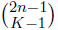 with the total number of classes 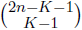. As a consequence, we only consider models for which 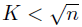 in the remainder.

##### *Remark* 3.2.

This formula was already obtained in a different context in Steel (1992) (Proposition 1) and, with a slightly different formulation, in Semple and Steel (2003, Proposition 4.1.4). In these works, the authors are interested in counting the “*r*-states convex characters on a binary tree”. Under the no-homoplasy assumption, this number can be shown to be equal to 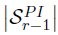.

#### 3.3 Another Characterization of Parsimony

The following proposition gives an alternative definition of parsimony under the no-homoplasy hypothesis using the linear formulation of the problem. It will be used for model selection in Section 4.2. Its technical proof is postponed to Appendix A.

##### Proposition 3.6

(Equivalence between parsimony and independence). *Let* **m**_**Y**_ *be a given mean vector*, **m**_**Y**_ ∈ ℝ^*n*^, *and* **Δ** *a vector of shifts such that* **TΔ** = **m**_**Y**_, *with* **T** *the tree matrix defined in Section 2.3*. *Under the no homoplasy assumption*, *the vector of shifts* **Δ** *is parsimonious if and only if the corresponding column-vectors of the tree matrix* 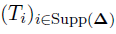 *are linearly independent*.

## 4 Statistical Inference

### 4.1 Expectation Maximization

Principle. As shown in Section 2.2, both BM and OUsun models can be seen as incomplete data models. The Expectation Maximization algorithm (EM, Dempster et al., 1977) is a widely used algorithm for likelihood maximization of these kinds of models. It is based on the decomposition: 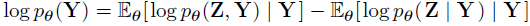. Given an estimate ***θ***^(*h*)^ of the parameters, we need to compute some moments of *p_**θ**^(h)^_*(**Z** | **Y**) (E step), and then find a new estimate 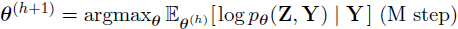. The parameters are given for the BM and OUsun in subsection 2.1. We assume here that the number of shifts *K* is fixed. We only provide the main steps of the EM. Additional details can be found in Appendix D.

**E step**. As **X** is Gaussian, the law of the hidden variables **Z** knowing the observed variables **Y** is entirely defined by its expectation and variance-covariance matrix, and can be computed using classical formulas for Gaussian conditioning. The needed moments of **Z** | **Y** can also be computed using a procedure that is linear in the number of tips (called “Upward-downward”) that takes advantage of the tree structure and bypasses inversion of the variance-covariance matrix (see Lartillot, 2014, for a similar algorithm).

**Complete Likelihood Computation**. Using the model described in Section 2.2, we can use the following decomposition of the complete likelihood: 
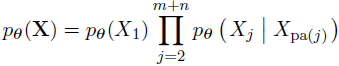

Each term of this product is then known, and we easily get 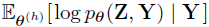.

**M step**. The difficulty comes here from the discrete variables (location of shifts on the branches). The maximization is exact for the BM but we only raise the objective function for the OUsun, hence computing a Generalized EM (GEM, seeDempster et al., 1977). This stems from the independent increment nature of the BM: shifts only affect *p*_**θ**_ (*X_j_* | *X*_pa(*j*)_)) on the branches where they occur and the maximization reduces to finding the K highest components of a vector, which has complexity *O*(*n* + *K* log(*n*)). By contrast, OUsun has autocorrelated increments: shifts affect *p*_**θ**_ (*X_j_* | *X*_pa(*j*)_) on the branches where they occur and on all subsequent branches. Maximization is therefore akin to segmentation on a tree, which has complexity *O*(*n*^*K*^).

**Initialization**. Initialization is always a crucial step when using an EM algorithm. Here, we use the linear formulation 2.4 or 2.6, and initialize the vector of shifts using a Lasso regression. The selection strength α is initialized using pairs of tips likely to be in the same group.

### 4.2 Model Selection

**Model Selection in the iid Case with Unknown Variance**. Model selection in a linear regression setting has received a lot of attention over the last few years. In Baraud et al. (2009), the authors developed a non-asymptotic method for model selection in the case where the errors are independent and identically distributed (iid), with an unknown variance. In the following, we first recall their main results, and then adapt it to our setting of non-independent errors.

We assume that we have the following model of *independent* observations: 
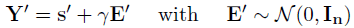
and we define a collection 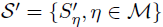 of linear subspaces of ℝ^*n*^ that we call *models,* and that are indexed by a finite or countable set ℳ. For each η ∈ ℳ, we denote by 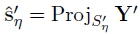 the orthogonal projection of **Y**′ on 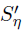, that is a least-square estimator of s′, and 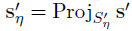 the projection of s′.

We extract from Baraud et al. (2009) the following theorem, that bounds the risk of the selected estimator, and provides us with a non-asymptotic guarantee. It relies on a penalty depending on the EDkhi function, as defined below:

#### Definition 4.1

(Baraud et al. (2009), Section 4, definitions 2 and 3). Let *D*, *N* be two positive integers, and *X_D_*, *X_N_* be two independent χ^2^ random variables with degrees of freedom *D* and *N* respectively. For *x* ≤ 0, define 
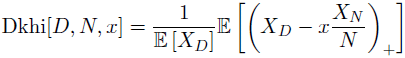

And define EDkhi[*D*,*N*, *q*] as the unique solution of the equation Dkhi[*D*,*N*, EDkhi[*D*,*N*, *q*]] = *q* (for 0 < *q* ≤ 1).

#### Theorem 4.1

(Baraud et al. (2009), Section 4, theorem 2 and corollary 1). *In the setting defined above*, *let D*_*η*_ *be the dimension of* 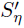, *and assume that N*_*η*_ = *n* − *D*_*η*_ ≥ 2 *for all η* ∈ ℳ. *Let* 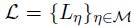 *be some family of positive numbers such that* 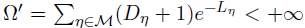, *and assume that, for A > 1*, 
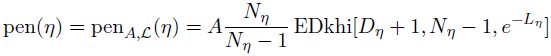

*Take* 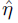 *as the minimizer of the criterion:* 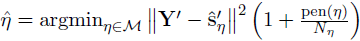. *Then*, *assuming that N*_*η*_ ≥ 7 *and* max(*L*_*η*_, *D*_*η*_) ≤ *κn for any η* ∈ ℳ, *with κ* < 1, *the following non-asymptotic bound holds:* 
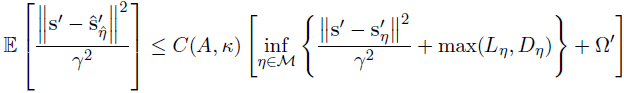

The penalty used here ensures an oracle inequality: in expectation, the risk of the selected estimator is bounded by the risk of the best possible estimator of the collection of models, up to a multiplicative constant, and a residual term that depends on the dimension of the oracle model. Note that if the collection of models is poor, such an inequality has low value. We refer to Baraud et al. (2009) for a more detailed discussion of this result.

**Adaptation to the Tree-Structured Framework**. We use the linear formulation described in 2.3, and assume that we are in the OUsun model (this procedure would also work for a BM with a deterministic root). Then, if **V** is a matrix of size *n*, with 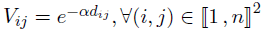, we have: 
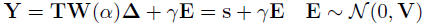

We assume that α is fixed, so that the design matrix **TW**(*α*) and the structure matrix **V** are known and fixed. A *model* is defined here by the position of the shifts on the branches of the tree, i. e. by the non-zero components of **Δ** (with the constraint that the first component, the root, is always included in the model). We denote by 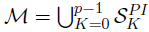 the set of allowed (parsimonious) allocations of shifts on branches (see Section 3.2), *p* being the maximum allowed dimension of a model. From proposition 3.6, for *η* ∈ ℳ, the columns vectors **T**_*η*_ are linearly independent, and the model *S*_*η*_ = Span(**T**_**i**_, *i* ∈ *η*) is a linear sub-space of ℝ^*n*^ of dimension D_*η*_ = |*η*| = *K*_*η*_ + 1, *K*_*η*_ being the number of shifts in model *η*. Note that as **W**(*α*) is diagonal invertible, it does not affect the definition of the linear subspaces. The set of models is then 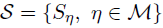.

We define the Mahalanobis norm associated to **V**^−1^ by: ‖**R**‖_**v**^−1^_ = **R**^***T***^V ^−1^**R**, ∀**R** ∈ ℝ^*n*^. The projection on S_n_ according to the metric defined by V^−1^ is then: 
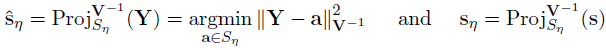

For a given number of shifts *K*, we define the best model with *K* shifts as the one maximizing the likelihood, or, equivalently, minimizing the least-square criterion for models with *K* shifts: 
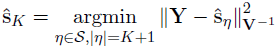

The idea is then to slice the collection of models by the number of shifts *K* they employ. Thanks to the EM algorithm above, we are able to select the best model in such a set. The problem is then to select a reasonable number of shifts. To compensate the increase in the likelihood due to over-fitting, using the model selection procedure described above, we select *K* using the following penalized criterion: 
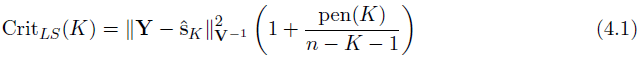

As noted in Baraud et al. (2009), the previous criterion can equivalently be re-written in term of likelihood, as: 
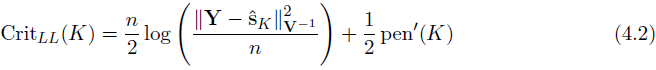
with 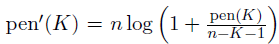. As we use maximum-likelihood estimators, we chose this formulation for the implementation. The following proposition then holds:

#### Proposition 4.1

(Form of the Penalty and guaranties (*α* known)). *Let* 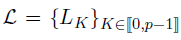, *with* 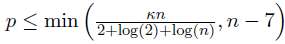, *the maximum dimension of a model*, *with κ* < 1, *and:* 
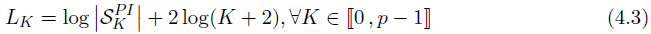

*Let* A > 1 *and assume that* 
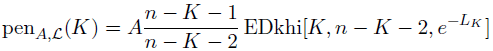

*Suppose that 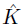 is a minimizer of 4.1 or 4.2 with this penalty*. *Then:* 
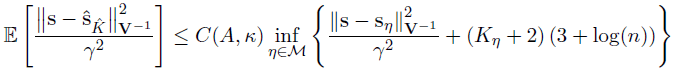
*with C*(*A*, *k*) *a constant depending on A* and *κ only*.

The proof of this proposition can be found in Appendix E. It relies on theorem 4.1, adapting it to our tree-structured observations.

#### *Remark* 4.1.

With this oracle inequality, we can see that we are missing the oracle by a log(*n*) term. This term is known to be unavoidable, see Baraud et al. (2009) for further explanations.

#### *Remark* 4.2.

Note that the chosen penalty may depend on the topology of the tree through the term 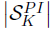 (see Section 3.2).

#### *Remark* 4.3.

The penalty involves a constant *A* > 1, that needs to be chosen by the user. Following Baraud et al. (2009) who tested a series of values, we fixed this constant to *A* = 1.1.

## 5 Simulations Studies

### 5.1 Simulations Scheme

We tested our algorithm on data simulated according to an OUsun, with varying parameters. The simulation scheme is inspired by the work of Uyeda and Harmon (2014). We first generated three distinct trees with, respectively, 64,128 and 256 tips, using a pure birth process with birth rate λ = 0.1. The tree heights were scaled to one, and their topology and branch lengths were fixed for the rest of the simulations. We then used a star-like simulation study scheme, fixing a base scenario, and exploring the space of parameters one direction at the time. The base scenario was taken to be relatively “easy”, with *β*_1_ = 0 (this parameter was fixed for the rest of the simulations), *α_b_* = 3 (i.e *t*_1/2,b_ = 23%), 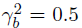 and *K_b_* = 5. The parameters then varied in the following ranges: the phylogenetic half life *t_1/2_* = ln(2)/*α* took 11 values in [0.01, 10]; the root variance 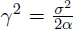 took 9 values in [0.05, 25]; the number of shifts *K* took 9 values in [0, 16] (see Figures 6–7 for the exact values taken). The problem was all the more *difficult* that *γ*^2^, *t*_1/2_ or *K* were large.

For each simulation, the *K* shifts were generated in the following way. First, their values were drawn according to a mixture of two Gaussian distributions, 𝒩(4,1) and 𝒩(−4,1), in equal proportions. The mixture was chosen to avoid too many shifts of small amplitude. Then, their positions were chosen to be balanced: we first divided the tree in *K* segments of equal heights, and then randomly drew in each segment an edge where to place a shift. We only kept parsimonious allocations.

Each of these configurations was repeated 200 times, leading to 16200 simulated data sets. An instance of a tree with the generated data is plotted in Figure 5.

**Figure 5:**
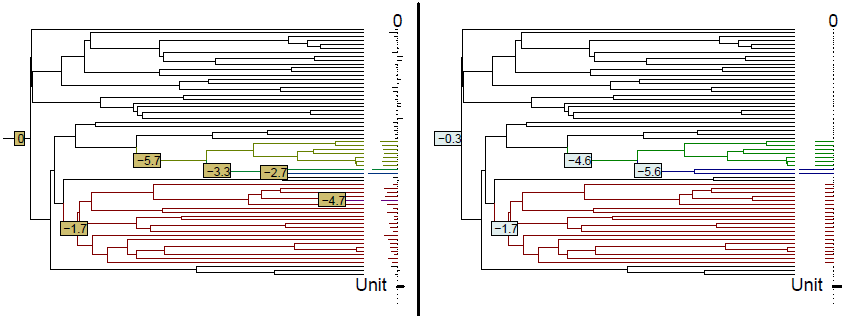
*Left*: Simulated configuration (with *t*_1/2_ = 0.75, *γ*^2^ = 0.5 and *K* = 5). The shifts positions and values are marked on the tree. The value of the character generated (positive or negative) is represented on the right. The colors of the branches correspond to the true regimes, black being the ancestral state. *Right*: One of the three equivalent allocations of shifts for the model inferred from the data, with corresponding vector of mean tip values. Shifts not recovered are located on pendant edges, and have low influence on the data. The two other equivalent allocations can be easily deducted from this one.

### 5.2 Inference Procedures

For each generated dataset, we ran our EM procedure with fixed values of 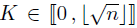, *n* being the number of tips of the tree. Remark that for *n* = 64, 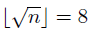, and we have no hope of detecting true values of *K* above 8 (see remark 3.1 for an explanation of the bound in 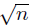). The number of shifts *K_s_* was chosen thanks to our penalized criterion, and we kept inferences corresponding to both *K*_s_ and the true number *K_t_*.

We ran two sets of estimations for *α* either known or estimated. The computations took respectively 66 and 570 (cumulated) days of CPU time. This amounts to a mean computational time of around 6 minutes (367 seconds) for one estimation when *α* is fixed, and 52 minutes (3137 seconds) when *α* is estimated, with large differences between easy and difficult scenarios.

### 5.3 Scores Used

The convergence of the EM algorithm was assessed through the comparison of the likelihood of the true and estimated parameters, and the comparison of mean number of EM steps needed when *α* is fixed or estimated. The quality of the estimates of *β*_1_, *t*_1/2_ and γ^2^ was assessed using the coefficient of variation. The model selection procedure was evaluated by comparing the true number of shifts with the estimated one, which should be lower. We do not expect to find the exact number as some shifts, which are too small or too close of the tips, cannot be detected. To evaluate the quality of the clustering of the tips, the only quantity we can observe, we used the Adjusted Rand Index (ARI, Hubert and Arabie, 1985) between the true clustering of the tips, and the one induced by the estimated shifts. The ARI is proportional to the number of concordant pairs in two clusterings and has maximum value of 1 (for identical clusterings) and expected value of 0 (for random clusterings). Note that this score is conservative as shifts of small intensity, which are left aside by our model selection procedure, produce “artificial” groups that cannot be reconstructed.

### 5.4 Results

The selection strength is notoriously difficult to estimate, with large ranges of values giving similar behaviors (see Thomas et al., 2014). We hence first analyse the impact of estimating α on our estimations, showing that the main behavior of the algorithm stays the same. Then, we study the shifts reconstruction procedure.

**Convergence and Likelihood**. For *α* known, all estimations converged in less than 49 iterations, with a median number of 13 iterations. For *α* estimated, the number of iterations increased greatly, with a median of 69, and a fraction of estimations (around 3.3%) that reached the maximum allowed number (fixed at 1000 iterations) without converging. Unsurprisingly, the more difficult the problem, the more iterations were needed. The log-likelihoods of the estimated parameters are close to the true ones, even when α is estimated (see supplementary Figure 13 in Appendix F, first row).

**Estimation of continuous parameters**. Figure 6 (first row) shows that we tend to slightly over-estimate *α* in general. The estimation is particularly bad for large values of *α* (with a high variance on the result, see first box of the row), and low values of α. In this regime, the model is “over-confident”, as it finds a higher selection strength than the real one and therefore a smaller variance (second row of Figure 6). For smaller and bigger trees, the estimators behave in the same way, but with degraded or improved values, as expected. We also note that taking the true number of shifts instead of the estimated one slightly degrades our estimation of these parameters (see supplementary Figure 13 in Appendix F). The estimation of *β*_1_ is not affected by the knowledge of *α* or *K* (see Figure 7, first row), and only has an increased variance for more difficult configurations. In the remainder, we only show results obtained for estimated α as estimating *α* does not impact ARI, 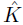 and 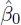 (see supplementary Figure 14 in Appendix F). the algorithm with the true number of shifts, γ^2^ is highly under-estimated. We will see afterwards that this is due to “undetectable” shifts.

**Estimation of the number of shifts**. The way shifts were drawn ensures that they are not too small in average, and that they are located all along the tree. Still, some shifts have a very small influence on the data, and are hence hard to detect (see Figure 5). The selection model procedure almost always under-estimates the number of shifts, except in very favorable cases (Figure 7, second row). This behavior is nonetheless expected, as allowing more shifts does not guarantee that the right shifts will be found (see supplementary Figures 11 and 12 in Appendix F).

**Figure 6:**
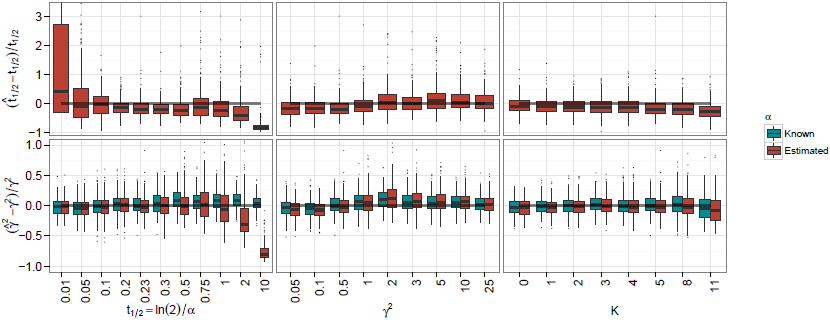
Box plots over the 200 repetitions of each set of parameters, for the phylogenetic halflife (top) and root variance (bottom) with *K* estimated, and *α* fixed to its true value (blue) or estimated (red), on a tree with 128 taxa. For better legibility, the y-axis of these two rows were re-scaled, omitting some outliers (respectively, for *t*_1/2_ and γ^2^, 0.82% and 0.46% of points are omitted). The whisker of the first box for *t*_1/2_ goes up to 7.5.

**Figure 7:**
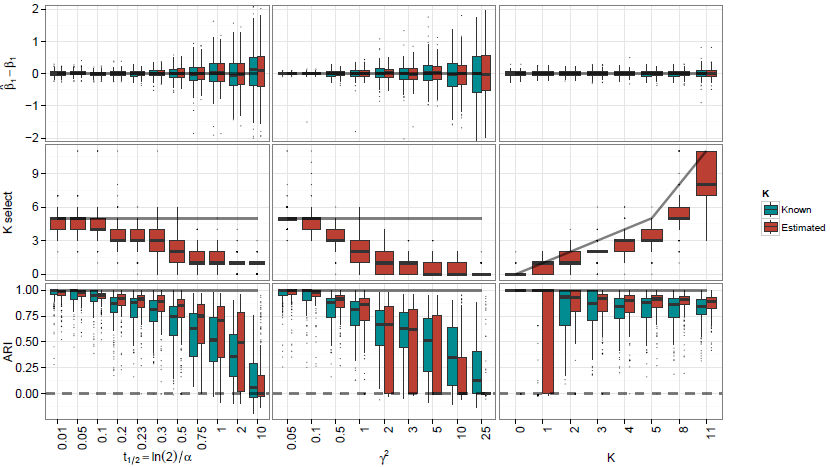
Same for *β*_1_(top), the number of shifts (middle) and ARI (bottom), with *α* estimated, and K fixed to its true value (blue) or estimated (red). As previously, the y-axis of the first row (*β*_1_) was re-scaled, omitting some outliers (1.38% of points are omitted).

**Clustering of the tips**. The ARI tends to be degraded for small values of *α* or high variance, but remains positive (Figure 7, third row). When only one shift occurs, the ARI is very unstable, but for any other value of *K*, it stays quite high. Finally, knowing the number of shifts does not improve the ARI.

**Equivalent Solutions**. When *α* and *K* are both estimated, only 5.8% of the configurations have 2 or more equivalent solutions. One inferred configuration with three equivalent solutions is presented Figure 5.

**Comparison with bayou**. As mentioned above, our simulation scheme, although not completely equivalent to the scheme used in Uyeda and Harmon (2014), is very similar, so that we can compare our results with theirs. The main differences lies in the facts that we took a grid on γ^2^ = *σ*^2^/2α instead of *σ*^2^, and that we took shifts with higher intensities, making the detection of shifts easier. We can see that we get the same qualitative behaviors for our estimators, with the selection strength *α* over or under estimated, respectively, in small or large values regions. The main difference lies in the estimation of the number of shifts. Maybe because of the priors they used (*K* ~ Conditional Poisson(λ= 9,*K_max_* = *n*/2)), they tend to estimate similar numbers of shifts (centered on 9) for any set of parameters. In particular, while our method seems to be quite good at detecting situations where there are no shifts at all, theirs seems unable to catch these kind of configurations, despite the fact that their shifts have low intensity, leading to a possible over-fitting of the data.

Overall, the behavior of the algorithm is quite satisfying. Our model selection procedure avoids over-fitting, while recovering the correct clustering structure of the data. It furthermore allows for a reasonable estimation of the continuous parameters, except for α which is notoriously difficult to estimate.

## 6 Case Study: Chelonian Carapace Lenxsgth Evolution

**Figure 8:**
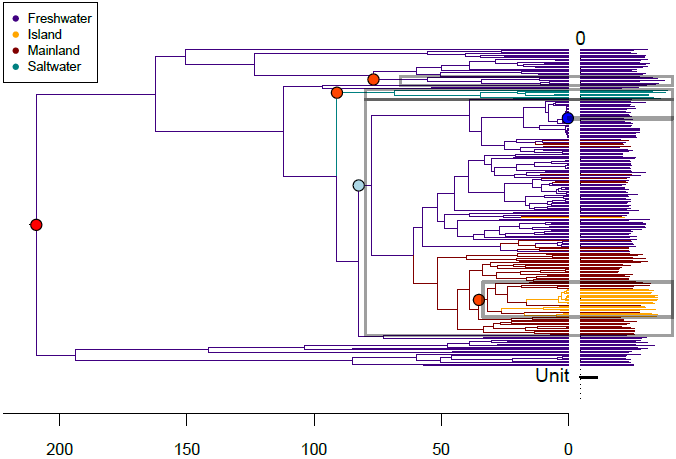
Phylogenetic tree of the Chelonians. Log-transformed traits values are represented on the right. Branch colors represent the habitats. The shifts found by our EM algorithm are shown as circles, with a color indicating the value of the shift, from blue (negative) to red (positive). Boxes highlight the groups induced by the shifts. The *x*-scale is in million years.

### 6.1 Description of The Dataset

Extant species of the order Testudines, or Chelonii, are turtles and tortoises, living all across the globe, and exhibiting a wide variation in body size, from the small desert speckled tortoise (*Homopus signatus*, 10 cm), to the large marine leatherback sea turtle (*Dermochelys coriacea*, 244 cm). In order to test the hypothesis of island and marine gigantism, that could explain the extreme variations observed, Jaffe et al. (2011) compiled a dataset containing a measure of the carapace length for 226 species, along with a phylogenetic tree of theses species, spanning 210 million years (my) (see Figure 8). They assigned each species to one of four habitats: mainland-terrestrial, freshwater, marine and island-terrestrial. Then, testing several fixed regimes allocations on the branches of the tree using the method described in Butler and King (2004), they found the best support in favor of a “OU2” model that assigned one regime to each habitat. Following Uyeda and Harmon (2014), we will refer to this model as the “OU_habitat_” model. Note that this model is ambiguously defined, as it requires to assign a habitat to each ancestral species. Using proposition 3.3, we found that there were 48 equivalent parsimonious ways of doing so that respect the habitats observed at the tips of the tree. One of these habitat reconstruction is presented Figure 8.

### 6.2 Method

We used the version of the dataset embedded in the package geiger (Harmon et al., 2008), that contains a phylogenetic tree and a vector of log-carapaces lengths. The corresponding habitats are reported in the Appendix of Jaffe et al. (2011).

We ran our algorithm with a number of shifts going from 0 to 20. Rather than estimating *α* directly within the EM as we did for the simulations, we took *α* varying on a grid, taking 6 values regularly spaced between 0.01 and 0.1, but fixed for each estimation. We found that this approach, although computationally more intensive, gave better results. These 6 × 20 = 120 estimations took around 2 hours of CPU time. For each number of shifts, going again from 0 to 20, we kept the solution with the maximal likelihood, and we applied the model selection criterion to them. This method gave a solution with 5 shifts, and a selection strength of 0.06 (i.e. 5.5 % of the total height of the tree). Using a finer grid for *α* gives highly similar results, allocating shifts to the same edges. These last estimations are given below.

### 6.3 Results

Our method selected a solution with 5 shifts, a rather strong selection strength (*t*_1/2_ = 5.4% of the tree height), and a rather low root variance (γ^2^ = 0.22, see table 1, first column). Two of those shifts are closely related to the habitats defined in Jaffe et al. (2011) (see Figure 8). The ancestral optimal value, that applies here to two clades of freshwater turtles, is estimated to be around 38 cm. A small decrease in size for a large number of mainland and freshwater turtles is found (optimal value 24 cm). Marine turtles (super-family Chelonioidea) are found to have an increased carapace length (with an optimal value of 130 cm), as well as a clade containing soft-shell tortes (family Trionychidae, optimal size 110 cm), and a clade containing almost all island tortoises, including several sub-species of Galápagos tortoises (*Geochelone nigra*). Only the Ryukyu black-breasted leaf turtle (*Geoemyda japonica*), endemic to the Ryukyu Islands in Japan, and distant on the phylogenetic tree, is not included in this group. Note that the group also contains some mainland tortoises of the genus Geochelone, that are closely related to Galaapagos tortoises. This is typical of our method: it constructs groups that are both phenotypically and phylogenetically coherent. Finally, one species is found to have its own group, the black-knobbed map turtle (*Graptemys nigrinoda*), with a very low optimal value of 1.4 × 10^−20^ cm, for a measured trait of 15 cm. The fact that the shift has a very high negative value (−49 in log scale) is just an artifact due to the actualization factor on a very small branch (0.18 my, for an inferred phylogenetic half-life of 11 my). This is a rather unexpected choice of shift location. When considering the linear model as transformed by the cholesky matrix of the variance to get independent errors (as in the proof of proposition 4.1), we find a leverage of 0. 94, indicating that this species trait behaves in the transformed space as an outsider.

### 6.4 Comparison with other Methods

In order to compare our results to previously published ones, we reproduced some of the analysis already conducted on this dataset. We hence ran the methods described in Jaffe et al. (2011) (using the `R` package `OUwie`, with fixed positions for the shifts), Uyeda and Harmon (2014) (implemented in package `bayou`), Ingram and Mahler (2013) (package `SURFACE`), and Ho and Ané (2014b) (function `OUshifts` in package `phylolm`). See Section F.3 in Appendix F for more details on these methods and the parameters we used. hours of CPU time.

The shifts allocated on the tree by methods bayou, `SURFACE` and `OUshifts` are presented on Supplementary Figure 15 (Appendix F). We can see that 3 among the most strongly supported shifts in the posterior distribution given by bayou, as well as some among the oldest shifts found by SURFACE and OUshifts are similar to the ones found by our method. The bayou method finds equal support for many shifts, all over the tree, and the median of the posterior distribution is 17 shifts, which is pretty close to the mode of the prior put on the number of shifts (15). The `SURFACE` and `OUshifts` methods select respectively 33 and 8 shifts, including many on pendant edges, that are not easily interpretable. The backward step of `SURFACE` allowed to merge the regimes found for marine turtles and soft-shell tortoises that our method found to have very similar optimal values. The results of the five methods are summarized Table 1. Note that these models are not nested, due to the status assigned to the root, and to the possible convergences.

Compared to step-wise heuristics, our integrated maximum likelihood based approach allows us to have a more “global” view of the tree, and hence to select a solution that accounts better for the global structure of the trait distribution. Thanks to its rigorous model selection procedure, our model seems to report significant shifts only, that are more easily interpretable than the solutions found by other methods, and that do not rely on any chosen prior.

**Table 1:** Summary of the results obtained with several methods for the Chelonian Dataset. For bayou, the median of the posterior distributions is given.

|  | Habitat | EM | bayou | SURFACE | OUshifts |
| --- | --- | --- | --- | --- | --- |
| Number of shifts | 16 | 5 | 17 | 33 | 8 |
| Number of regimes | 4 | 6 | 18 | 13 | 9 |
| lnL | -133.86 | -97.59 | -91.54 | 30.38 | -79.79 |
| Marginal lnL | NA | NA | -149.09 | NA | NA |
| $\alpha$ ($\times h$, per my) | 9.32 | 12.76 | 36.54 | 1.72 | 3.25 |
| $\ln 2/\alpha$ (my) | 15.56 | 11.36 | 3.97 | 84.28 | 44.64 |
| $\sigma^2$ ($\times h$, per my) | 6.21 | 5.57 | 11.91 | 0.72 | 2.29 |
| $\gamma^2$ | 0.33 | 0.22 | 0.16 | 0.21 | 0.35 |
| CPU time (min) | 65.25 | 134.49 | 136.81 | 634.16 | 8.28 |

## Acknowledgments

We would like to thank Cecile Ane for helpful discussions on an early draft of this manuscript. We are grateful to the INRA MIGALE bioinformatics platform (http://migale.jouy.inra.fr) for providing the computational resources needed for the experiments. We also thank the two anonymous reviewers whose careful and critical reading greatly helped improve this manuscript.

## A Technical Proofs of Section 3

### *Proof of Proposition 3.1*.

Let 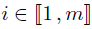 be an internal node with *L_i_* children nodes (*d*_1_, …, *d*_*L*_*i*__) and **K**^*i*^ the corresponding vector, defined as in the proposition. Then, for any 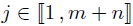: 
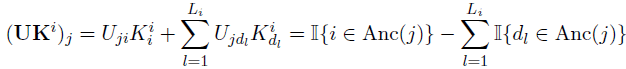

We can then distinguish three possibilities:

- If *i* ∉ Anc(*j*), then *d_l_* ∉ Anc(*j*) for all 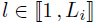 and (**TK**^*i*^)*j* = 0.
- If *j* = *i*, then *i* ∈ Anc(*j*), and, by definition, *d_l_* ∉ Anc(*j*) for any 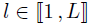, so (**UK**^*i*^)_*i*_ = 1.
- Else, if *i* is an ancestor of *j*, with *i* ≠ *j*, then, as *i* is internal, one (and only one) of its child *d_l_* is also an ancestor of *j* (potentially *j* itself), so that the sum cancels out.

This proves that 
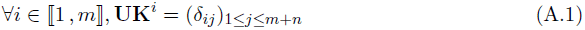

In particular, as **TK**^*i*^ is the vector of the last *n* coordinates of **UK**^*i*^, this shows that the vectors (**K**^1^, …, **K**^*m*^) are in the kernel of **T**.

Then, as we found *m* independent vectors in the kernel of **T**, which is a space of dimension lower than *m* (as the *n* columns of **T** representing tips are linearly independent, and by the rank theorem), this family of vector is a basis of the kernel space.

### Proof of lemma 3.1.

First, (**b**_*m*+1_, …, **b**_*m*+*n*_) is a family of *n* independent vectors of *S* of dimension *n*, so is a basis of *S*, and *b*′ is a basis adapted to ker(**T**) ⨁ *S*.

Let 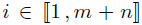. Let’s show that 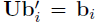. If *m* + 1 ≤ *i* ≤ *m* + *n*, then 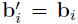, and **Ub**_*i*_ = **b**_*i*_ is the *i^th^* column of **U**. Otherwise, if 1 ≤ *i* ≤ *m*, then 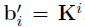, and, from equation A.1, 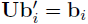. This shows that **U** is the change of basis matrix between *b* and *b*′.

### Proof of Proposition 3.6.

By contraposition, let’s first assume that the vector-columns 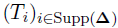 are linearly dependent, and prove that **Δ** is not parsimonious. This means that we can find a vector **E**, **E** ∈ ℝ^*m*+n^, such that Supp(**E**) ⊂ Supp(**Δ**), and **TE** = 0. We can hence find *j* ∈ Supp(Δ), *j* > 1, such that *E_j_* ≠ 0. Then if λ = − **Δ**_j_/*E_j_*, the vector **Δ**′ = **Δ** + λE is a vector of shifts on the tree with one less non-zero coordinate than **Δ** such that **TΔ**′ = **m_Y_**. Hence, **Δ** is not parsimonious.

Reciprocally, by contraposition, assume that **Δ** is not parsimonious. Then by proposition 3.4, it produces *p* groups, with *p* ≤ *K*. Hence the application associated with 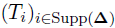 goes from a space of dimension *K* +1 to a space of dimension *p* ≤ *K*, and hence is not injective, and the family 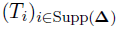 is not independent.

## B Enumeration Of Equivalence Classes

### Definition B.1

(Coloring concatenation). Let *i* be a node of tree 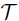 with *L_i_* daughter nodes (*i*_1_, …, *i*_*L_i_*_), *L_i_* ≥ 2, and assume that the tips are colored according to the application 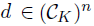. We denote by 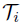 the sub-tree rooted at node *i*, and by 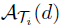 the set of parsimonious shifts allocations on 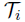 that produce a coloring of the tips compatible with *d*. At the root, 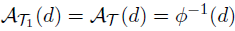.

- For 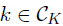, *S_i_*(*k*) is the *cost* of starting from node *i* with color *k*, *i.e.* the minimal number of shifts needed to get the right coloring of the tips of 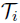, when starting with color *k*.
- 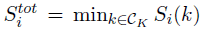 is the minimal cost of subtree 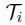, *i.e.* the number of shifts of a parsimonious coloring. 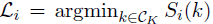 is the set of colors root *i* can take in a parsimonious coloring of sub-tree 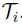.
- For 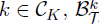 is the set of colorings of 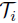 that respect the colors at the tips, have *S_i_*(*k*) shift, and start with color *k*.
- For 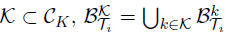. Hence, 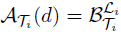, and the computation of 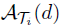 only requires the computation of *S_i_*(*k*) and 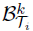 for any 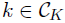.
- For 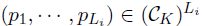, and 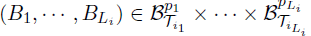 we define, for 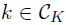, the concatenation 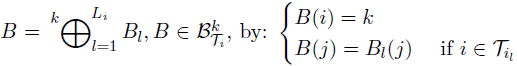. As the sub-trees 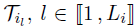 do not overlap, this application is correctly defined on the nodes of 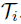.

Using these definitions, we can state the following recursion formula:

### Proposition B.1

(Enumeration Recursion Formula). *Let* 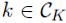, and 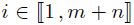. *If i is a tip of the tree*, *then* 
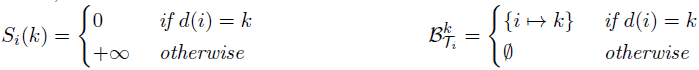

*If i is a node of tree* 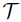 *with L_i_ daughter nodes* (*i*_1_, …, *i*_*L_i_*_), *L_i_* ≥ 2, *and assuming that S_il_*(*k*) *and* 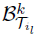 *are known for any* 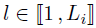 *and* 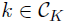, *define*, *for* 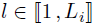: 
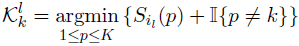

*As these sets are not empty*, *let* 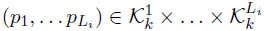. *Then* 
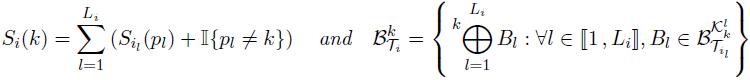

### Proof.

The actualization of *S_i_*(*k*) is the same as in the Sankoff algorithm (Sankoff, 1975). The set 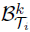 is then obtained by enumerating all the possible ways of concatenating children sets 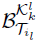, each of which is the ensemble of solutions for the sub-tree 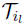 that realize the minimal number of shifts when starting in state *k*.

Remarking that 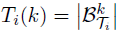, proposition 3.3 of the main text follows immediately.

## C A Vandermonde Like Identity

### Proposition C.1.

*Let* (*n*,*n*′) ∈ *N and K* ∈ ℕ. *With the standard convention that* 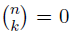 *if n* < *k*, 
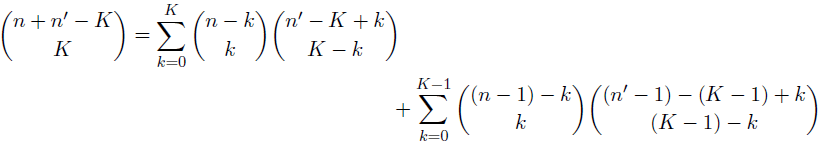
*which can be rewritten in a more symmetric way as:* 
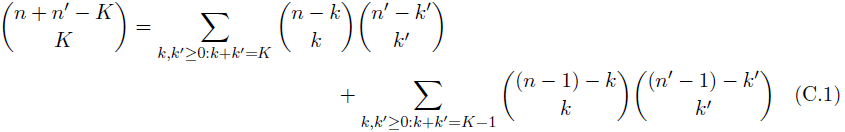
*Similarly*, 
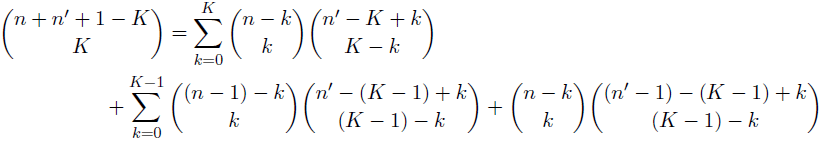
*which can be rewritten in a more symmetric way as:* 
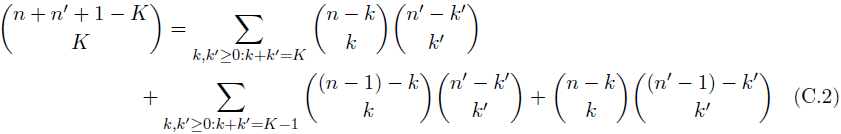

Note that Eq (C.1) generalizes in some way the Vandermonde identity which states 
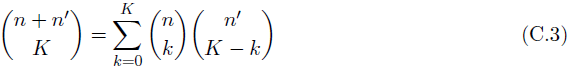

Although several proofs of the Vandermonde identity are known (geometric, algebraic and combinatorial), we only provide a geometric proof of this Vandermonde-like identity.

Consider a grid of size (*n* + *n*′) × *K*. We are interested in grid-valued paths that can move either by (1, 0) or by (2,1). In other words, if the *k*^th^ position of a path is (*x_k_*, *y_k_*), then its next position (*x*_k+1_,*y*_*k*+1_) is either (*x_k_* + 1,*y_k_*) or (*x_k_* + 2,*y_k_* + 1). We are interested in paths starting at (0,0) and ending at (*n* + *n*′, *K*).

Such a path consists of *K* moves of type (2,1) and *n* + *n*′ − 2K moves of type (1,0) and is uniquely determined by the positions of the moves of the former type. There are 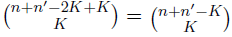 distinct positions and therefore as many such paths.

We now sort the paths according to the value *i* they take when either reaching the line *x* = *n* or reaching the line *x* = *n* + 1 without reaching the line *x* = *n* first. We refer to the latter paths as crossing the line *x* = *n*. Note that this sorting induces a partition of all paths (see Figure 9)

**Figure 9:**
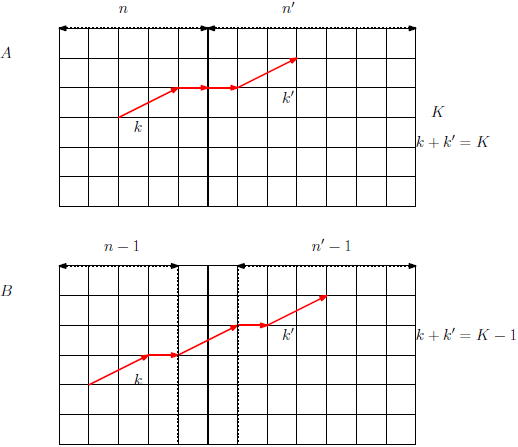
Partition of paths according to whether they reach (A) or cross (B) the line *x* = *n*

A path reaching *x* = *n* at position *i* uniquely gives rise to two paths: one from (0, 0) to (*n*, *i*) and one from (*n*, *i*) to (*n* + *n*′, *K*) or equivalently from 0 to (*n*′, *K* − *i*). There are 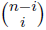 different paths of the first kind and 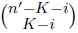 of the second. There are therefore 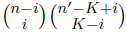 paths that pass through (*n*,*i*).

A path crossing the line *x* = *n* and reaching the line *x* = *n* + 1 at *i* must do so with a last move of type (2,1). It therefore uniquely defines a path from (0,0) to (*n* − 1, *i* − 1) and a path from (*n* + 1, *i*) to (*n* + *n*′, *K*), or equivalently from (0,0) to (*n*′ − 1, *K* − *i*). There are therefore 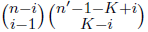 paths that cross the line *x* = *n* and pass through (*n* + 1, *i*).

Putting everything together, we get: 
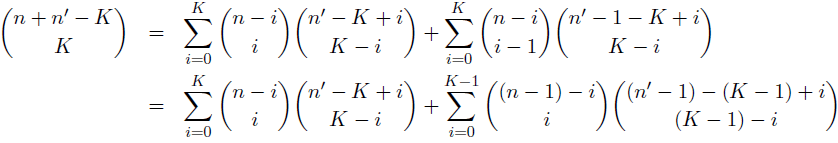
which is exactly Eq. (C.1).

To prove Eq. (C.2), we start from a grid of size (*n* + *n*′ + 1) × *K* and are again interested in the paths starting from the bottom left corner and ending in the upper right corner using only (2, 1) and (1, 0) moves. These paths have exactly *K* moves of type (2, 1) and there are 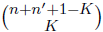 of them. This time, we partition paths upon the move observed between *x* = *n* and *x* = (*n* + 1).

The move can be (see also Figure 10):

- (1, 0), in which case *k* (resp.*k*′) moves of type (2,1) are used in the interval [1,n] (resp. [*n* + 1, *n* + *n*′1]) such that *k* + *k*′ = *K*;
- (2, 1) starting from *x* = *n* and therefore ending at *x* = *n* + 2, in which case *k* (resp. *k*′) moves of type (2,1) are used in the interval [1,*n*] (resp. [*n* + 2,*n* + *n*′1]) such that *k* + *k*′ = *K* − 1 (one move (2, 1) has already been consumed);
- (2, 1) ending at *x* = *n* + 1 and therefore starting from *x* = *n* − 1, in which case *k* (resp. *k*′) moves of type (2,1) are used in the interval [1,*n* − 1] (resp. [*n* + 1,*n* + *n*′1]) such that *k* + *k*′ = *K* − 1 (one move (2, 1) has already been consumed);

**Figure 10:**
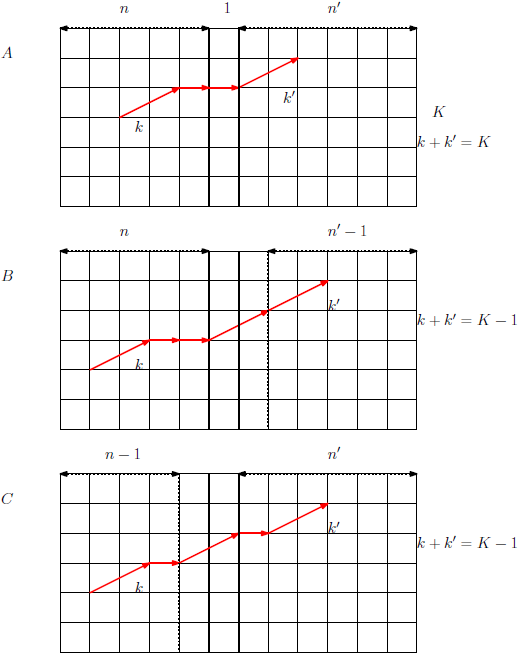
Partition of paths according to whether to the move used between *x* = *n* and *x* = *n* + 1. Cases A, B and C correspond to the items listed in the main text.

Wrapping everything together and using the same arguments as before, we get Eq. (C.2).

## D Technical Details of The EM

### D.1 E Step

Given a set of parameters **θ**^(*h*)^, we have: 
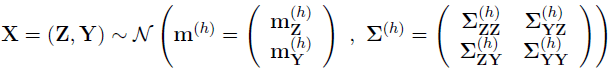
hence: 
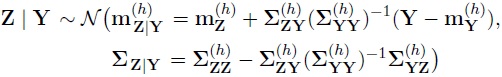

#### *Remark* D.1.

We can see that this approach forces us to invert 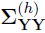, a *n* × *n* matrix, which is a costly operation, of order *O*(*n*^3^). It also computes the complete matrix ∑_**Z**|**Y**_ whereas we only need a linear number of its coefficients: conditional variances and covariances of the form ℂov [**Z**_*i*_; **Z**_pa(*i*)_ | Y]. Due to the tree structure and to the Gaussian nature of the processes studied, it is possible to compute all the quantities needed in a linear time, using a “forward-backward”-like algorithm (here, “upward-downward”, see Lartillot (2014) for a similar algorithm.). The upward step is similar to the pruning algorithm described in Felsenstein (2004, chap. 23). See also Ho and Ané (2014a) for an algorithm linear in the number of iterations.

### D.2 Complete Likelihood Computation

Using the incomplete data model described in Section 2.2, we can write: 
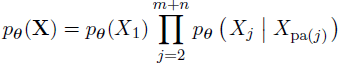

Taking the expectation, we get for the BM: 
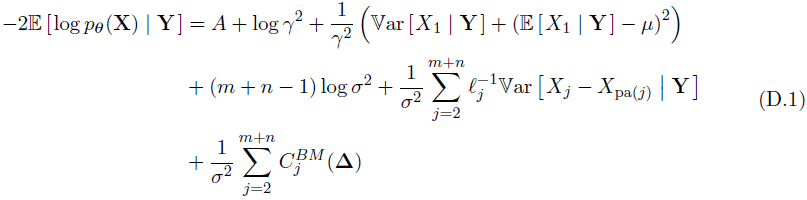
and, for the OUsun: 
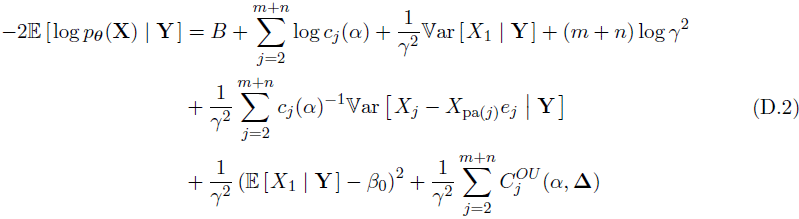
where *A* and *B* are constants, and for each node *j*, 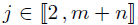, we define an actualization factor 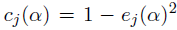, with 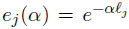, and 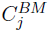 and 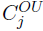 are *costs* associated with branch *b_j_*: 
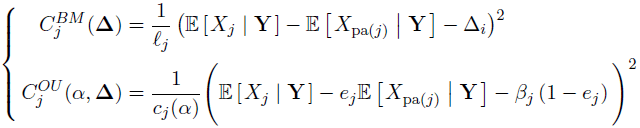

### D.3 M step

Assuming that *p*_***θ**^(*h*)^*_(**Z** | **Y**) is known, we need to compute ***θ***^(*h*+1)^ by maximizing 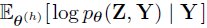. We have to deal with parameters of different nature, discrete or continuous. For a given vector **Δ**^(*h*+1)^ of *K* non-zero shifts, we can exhibit closed formulas for *μ*;^(*h*+1)^, *σ*^(h+1)^ and γ^(h+1)^, for the BM: 
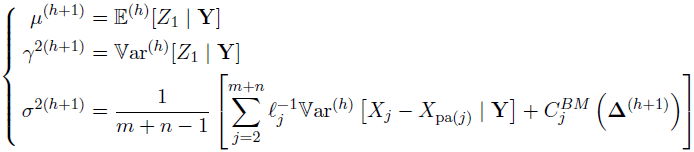
and, for the OUsun: 
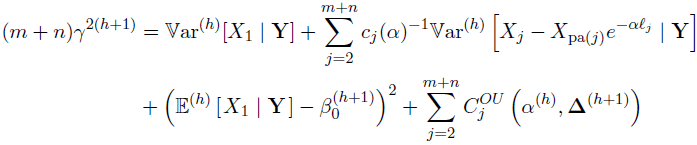

There is no such closed formula for α. In the implementation we propose, this parameter is actualized after all the others, by doing a numerical maximization of the objective function.

Finally, the vector **Δ**^(*h*+1)^ of *K* non-zero shifts can be chosen in an optimal way for the BM thanks to a simple algorithm explained below. In the OUsun case, we can only increase the objective function, and not maximize it. In that case, we hence use a Generalized EM algorithm (GEM, see Dempster et al., 1977).

**Optimal Shift Location for the BM**. We want to minimize the sum of costs: 
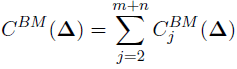

Each cost is associated to a branch *b_j_*, 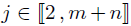, and, when the sum is minimal, 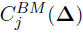 can only take two values: 
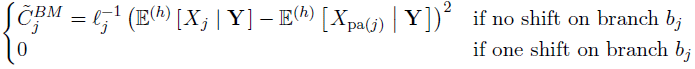

The sum can hence be minimized in the following way:

1. Compute 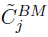 for all 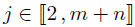.
2. Find the *K* highest costs 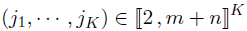.
3. Set 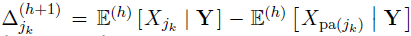 for all 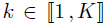, and 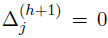 if *j* ≠ {*j*_l_, …, *j_K_*}.

This exact and fast algorithm works for the BM because all the costs are independent. Note that it would work for any Levy Process without memory, such as those proposed in Landis et al. (2013) to model evolution of quantitative traits.

**GM Step for Shifts Locations for the OU**. With *α^(h)^* fixed, we want to minimize the sum of costs: 
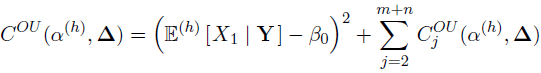

The previous algorithm does not work, because the costs are not independent. Solving the problem exactly would require to visit all the possible configurations, and the complexity would be too high, of order 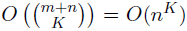. To reduce the execution time of the algorithm, we use heuristics to lower, if not minimize, the sum of costs. We use the following formulation: 
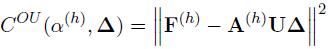
where **U** the complete tree matrix given in subsection 2.3, 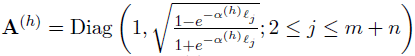 a diagonal matrix depending on *α*^(*h*)^, and **F**^(*h*)^ a vector of expectations, with 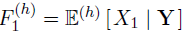, and, for 2 ≥ *j* ≥ *m* + *n*, 
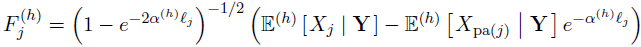

We can then use a Lasso algorithm to impose sparsity constraints on **Δ**. If **Δ**_−1_ is the vector of shifts without the initial value (intercept), then a Lasso estimator is given by, for A ≤ 0: 
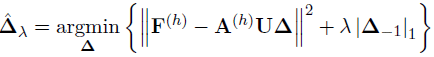

The estimated vectors 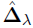 have a support that is sparser when λ becomes higher. One then only need to find the right penalty factor λ that ensure that the support has exactly *K* non zero coordinates, plus the initial value. We ensure that the *K* shifts are allocated in a parsimonious way by checking their linear independence, using proposition 3.6.

An other method is to take the previous solution **Δ**^(*h*)^, and test all the configurations where only one shift has moved, and take the best one. In both methods, one also has to ensure that the objective function is increased by the new choice of shifts, so that the GEM algorithm works correctly. This step is generally the longest one in one iteration of the EM.

### D.4 Initialization

Initialization is always a crucial step when using an EM algorithm. The vector of shifts **Δ** is initialized thanks to a Lasso procedure. To do that, we use the linear formulation 2.4 or 2.6 of the main text, and we calibrate the penalty so that the initialization vector has a non zero first coordinate (initial value), and *K* other non-zero coordinates. The variance-covariance matrix is initialized with defaults parameters, and is taken into account thanks to a Cholesky decomposition.

We also initialize the selection strength *α*. We use the following property: if *Y_i_* and *Y_j_* are two tips in the same group, then, under an OUsun, 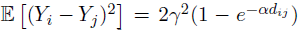. Using regression techniques, we can get an initial estimation of *α* and γ^2^ from all these couples. In practice, we first initialize the position of the shifts, and then use only pairs of tips from the same estimated group. Then, as the groups are only approximated, some of the selected pairs (*Y_i_*, *Y_j_*) might not share the same expectation, and we use a robust regression to get more accurate initial estimates.

## E Proof of Proposition 4.1 for Model Selection

We prove the proposition using the linear formulation 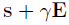, as derived in the main text for the OUsun (with s = **TW**(*α*)Δ, *γ*^2^ = *σ*^2/(2α)^, and *V_ij_* = *e*^−*α d_ij_*^, see Formula (2.5)). Note that this framework also holds for the BM with a fixed root (with *s* = **TΔ**, *γ* = a, and *V_ij_* = *t_ij_*, see Formula (2.4)).

We first handle the case where there are no correlations (**V** diagonal), and then use a Cholesky decomposition to handle the general case. Note that the case **V** diagonal can be seen as the limit of the OUsun when *α* = +∞, or as a BM on a star tree.

### *Case* **V** *Diagonal*

In the iid case, we just need to check the conditions of theorem 4.1. This paragraph is highly inspired by the derivation of the bound for the detection of non-zero mean components exposed in Baraud et al. (2009) (sub-section 5.2). Assume that *D_η_* = *K_η_* + 1 ≤ *p* ≤ *n* − 7 for all *η* ∈ ℳ. The estimator is defined by 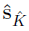, with: 
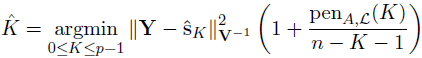

From the definition of 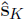, and as the penalty depends on the model only through its number of shifts, we get that 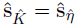 the minimizer of the criterion of theorem 4.1 (with *N_η_* = *n* − *D_η_* = *n* − *K_η_* − 1). We then have: 
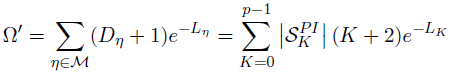

With the weights *L_K_* defined in equation 4.3 of the proposition, we get: 
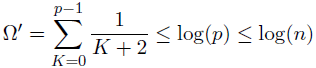

As: 
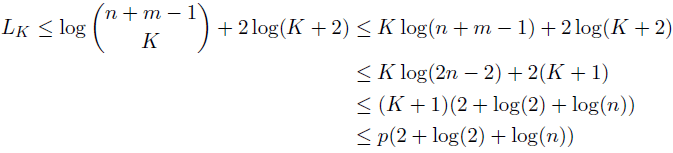
if 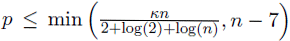, then max(*L_η_*,*D_η_*) ≥ *Kn* for any η ∈ ℳ, and we get the announced bound from the second proposition of theorem 4.1.

### *Case* **V** *not Diagonal*

Using a Cholesky decomposition, we can find a lower triangular matrix **L** such that **V** = **LL**^*T*^. Then, denoting **Y**′ = **L**^−1^**Y**, s′ = **L**^−1^s, and **E**′ = **L**^−1^**E**, we have **Y**′ = s′ + γ**E**′, with 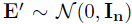, and we can apply theorem 4.1 as above. As we changed the metric, the estimators are projections on the linear spaces 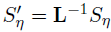 for *η* ∈ ℳ, and we have: 
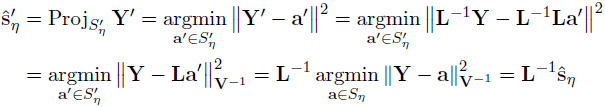

So 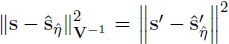 and 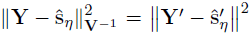, and, as the form of the penalty does not depend on *V*, by minimizing: 
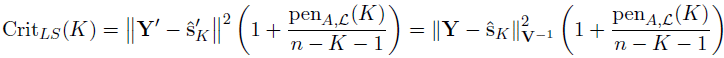
we get the announced bound on 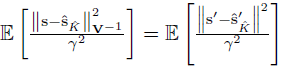.

## F Supplementary Figures

### F.1 Simulation Study: Sensitivity and False Positive Rate

**Definition of the Scores**. We denote by *TP* the number of True Positives, i.e. the predicted edges on which a shift actually occurred, and *FP* the number of False Positives. The sensitivity 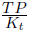 is the proportion of well predicted shifts among all shifts to be predicted, and the False Positive Rate (FPR) 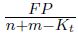 is the proportion of false positive among all edges with no shifts.

Note that here, due to the possible lack of identifiability, the position of the shifts on the tree is not well defined, as a shift can be on a particular edge for one of the equivalent solutions, but not on the others (see Section 3.1). These two scores are hence ill defined for our problem. To avoid such problems, we restrict ourselves to the 91% of unambiguous configurations that occurred during the simulations.

**Interpretation of Results**. Figure 11 shows that the FPR are systematically worse when using the true number of shifts, indicating that the additional shifts found when compared to the selected number are misplaced. The FPR remains very low, as only a small number of shifts is to be found. Unsurprisingly, the Sensitivity is on the contrary improved when taking the real number of shifts, as shown Figure 12. In addition, the sensitivity is highly degraded when *α* is small or γ^2^ is high, but does not exhibit a clear tendency in the real number of shifts, and the knowledge of the true value of α does not seem to matter.

**Figure 11:**
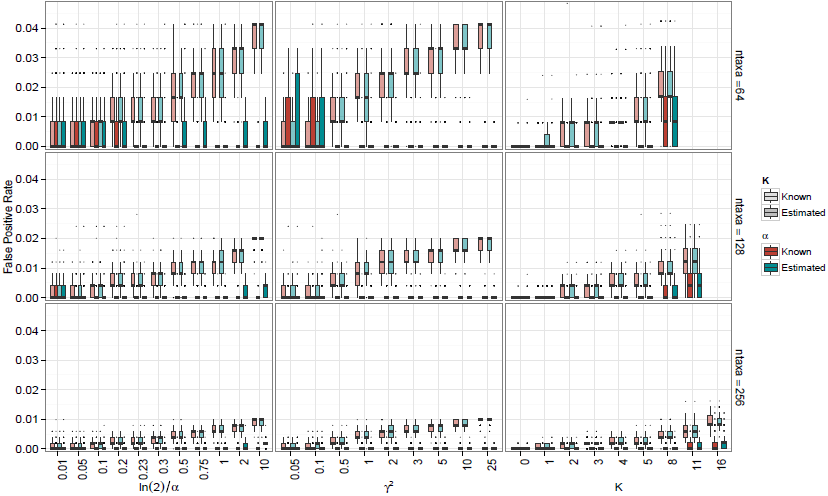
False Positive Rate computed for the different configurations. Note the *y* scale, that only goes to 0.05.

**Figure 12:**
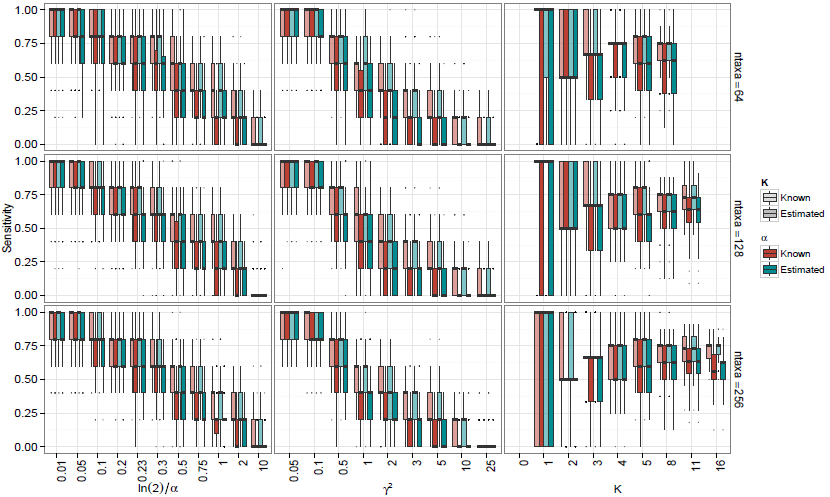
Sensitivity computed for the different configurations, with box-plots over the repetitions.

### F.2 Simulation Study: Complementary Analysis

To complete the analysis conducted in the main text, Figure 13 presents the variations of the log-likelihood, phylogenetic half-life, and root variance when *α* is estimated, and the number of shifts is known or estimated. We can see here that the likelihood is slightly higher when the number of shifts is fixed, which is coherent with the behavior of our model selection procedure, that tends to under-estimate the true number of shifts (see Figure 7 of the main text). We also note that knowing the true number of shifts has not a great influence on the estimation of *α* and γ, making the later worse, if anything.

**Figure 13:**
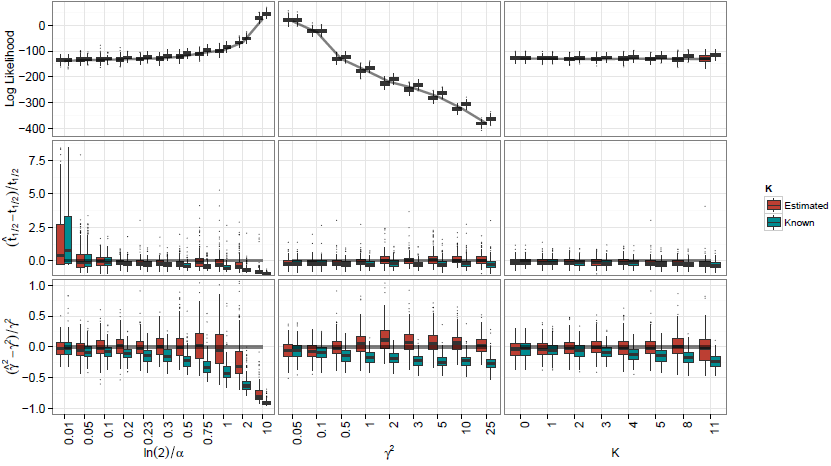
Box plots over the 200 repetitions of each set of parameters, for the log-likelihood (top), phylogenetic half-life (middle) and root variance (bottom) with *α* estimated, and *K* fixed or estimated, on a tree with 128 taxa.

Figure 14 shows the variations of the estimations of *β*_1_, the number of shifts and the ARI when the number of shifts is estimated, and α is fixed or estimated. This confirms our earlier statement, that not knowing α with precision does not have a great impact on the model selection procedure (see also Cressler et al., 2015).

**Figure 14:**
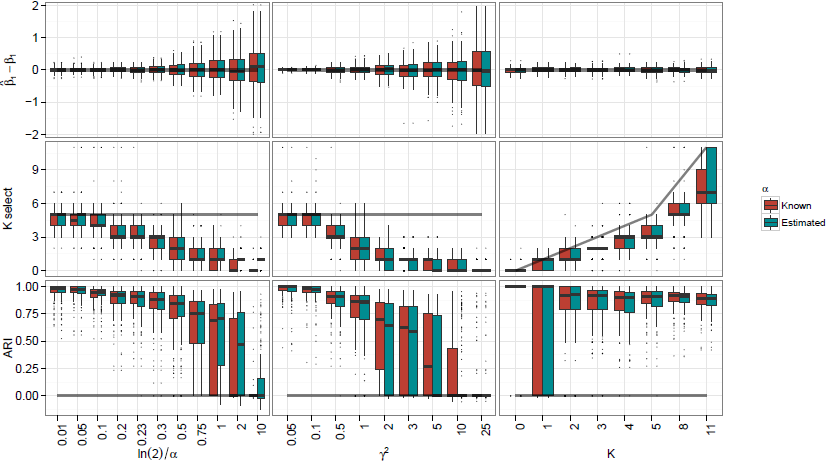
Same for *β*_1_ (top), the number of shifts (middle) and ARI (bottom), with *K* estimated, and a fixed or estimated.

### F.3 Chelonia Dataset: Comparison of Inferred Shift Locations

On Figure 15, we present and compare the shift locations found by our method, and methods bayou and `SURFACE`. The differences found are explored deeper in the main text (Section 6.4).

**Figure 15:**
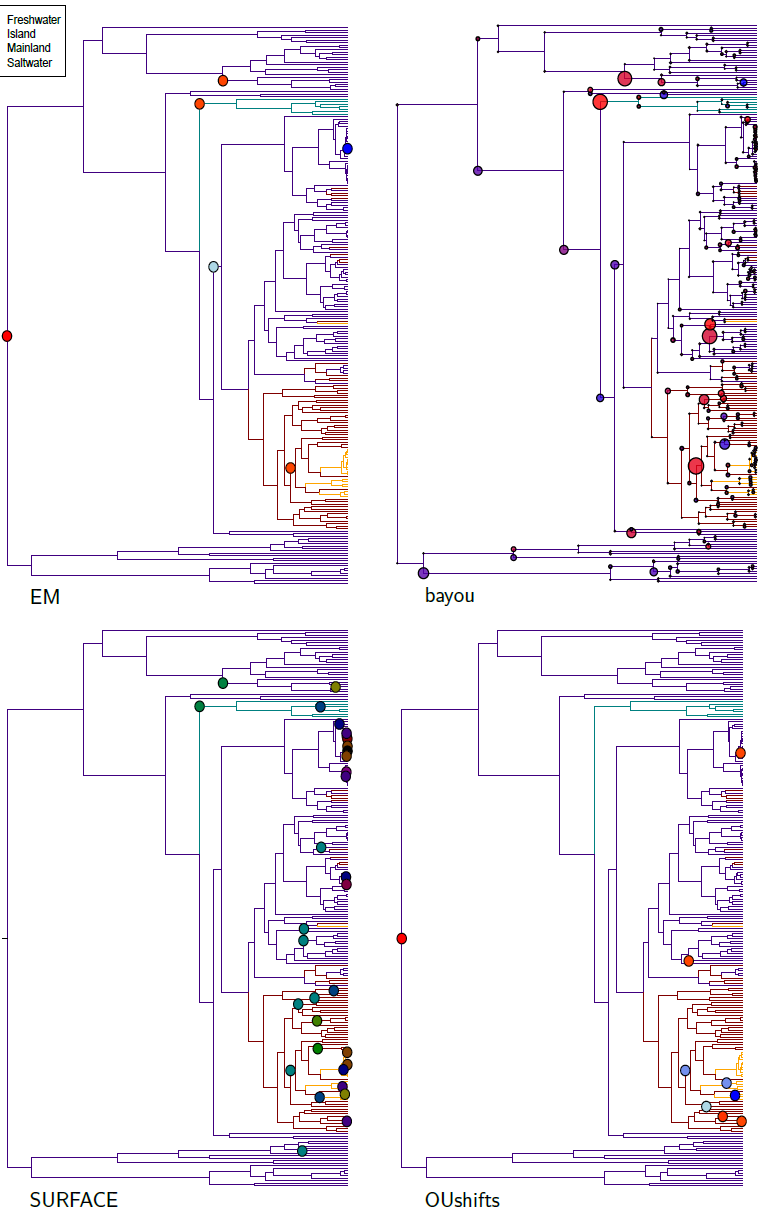
Solutions found by the EM (top left), bayou (top right), **SURFACE** (bottom left) and OUshifts (bottom right). The branch coloring represents the habitats. For the **EM**, bayou and OUshifts, the shifts coloring represents their values, from blue (negative) to red (positive). For bayou, the size of the circles are proportional to their posterior probability. For **SURFACE**, the 13 colors of the shifts represent the regimes.

**Details on the methods**. In this paragraph, we give more details on the 4 already existing methods that we compared to ours in the dataset. We first fitted an OU_habitat_ model with fixed regimes as in Jaffe et al. (2011), using the `R` package `OUwie` (Beaulieu et al., 2012). We tested all of the 48 possible ways of allocating internal nodes, and took the solution with the highest likelihood. Using the package `bayou` (Uyeda and Harmon, 2014), we reproduced the Bayesian analysis of the data, using two independent chains of 500000 generations each, discarding the first 150000 generations as burning. We assigned the priors that were used in the original study on the parameters, namely: *P*(α) ~ LogNormal(ln *μ* = −5, ln*σ* = 2.5), *P*(*σ*^2^) ~ LogNormal(ln *μ* = 0, ln*σ* = 2), *P*(*β*_1_) ~ Normal(*μ* = 3.5, *σ* = 1.5), *P*(*K*) ~ Conditional Poisson(λ = 15,*K_max_* = 113). The computations took around 2.3 hours of CPU time. We also ran the stepwise-AIC method `SURFACE` (Ingram and Mahler, 2013), that relies on a forward-backward procedure. This took around 11 hours of CPU time. Finally, we ran the function OUshifts from the package phylolm, that uses a modified BIC criterion and a heuristic stepwise procedure to detect shifts (Ho and Ané, 2014b). Thanks to an efficient linear algorithm, detailed in Ho and Ané (2014a), this function is pretty fast, taking only about 8 minutes of CPU time. Note that the model used in these last three methods (bayou, `SURFACE` and `OUshifts`) are slightly different from ours, as they assume that the root is fixed to the ancestral optimum state, and not drawn from its stationary distribution.

For better legibility, strips with *t*_1/2_, γ^2^ and *β*_1_ on these two figures were re-scaled, omitting some outliers (respectively, 0.21%, 0.27% and 0.27% of points are omitted). The whisker of the first box for *t*_1/2_ goes up to 7.5.

**Note on Computation Times**. We found that the running time for our method was similar to the running time of previous algorithms (see Table 1 in the main text). However, our computations can be highly parallelized, as each run for a fixed number of shift is independent from the others. For instance, in the previous example, the computation time could be divided by 6, each estimation for a fixed α running on a different core. On the contrary, the `SURFACE` method cannot be parallelized at all, and only independent chains can be parallelized for Bayesian methods, so that the computation time can only be divided by 2 in our example.

## G Practical Implementation

The statistical method described here was implemented on the statistical software `R` (R Core Team, 2014), and the code is freely available on GitHub (https://github.com/pbastide/Phylogenetic-EM). Phylogenetic trees were handled thanks to the package `ape` (Paradis et al., 2004). Packages `TreeSim` (Stadler, 2014), `robustbase` (Rousseeuw et al., 2014) and `quadrupen` (Grandvalet et al., 2012) were used, respectively, for random tree generation, robust regression and Lasso regression. The penalty described in proposition 4.1 is implemented in package `LINselect` (Baraud et al., 2013).

Parallelization was achieved thanks to `R` packages `foreach` (Weston, 2014b) and `doParallel` (Weston, 2014a).

Package `mclust` (Fraley et al., 2012) was used for **ARI** computations. Plots were made thanks to packages `ggplot2` (Wickham, 2009) and `reshape2` (Wickham, 2007).

